# Hybridization and the Coexistence of Species

**DOI:** 10.1101/2021.04.04.438369

**Authors:** Darren Irwin, Dolph Schluter

## Abstract

It is thought that two species can coexist if they use different resources present in the environment, yet this assumes that species are completely reproductively isolated. We model coexistence outcomes for two sympatric species that are ecologically differentiated but have incomplete reproductive isolation. The consequences of interbreeding depend crucially on hybrid fitness. When hybrid fitness is high, just a small rate of hybridization can lead to collapse of two species into one. Low hybrid fitness can cause population declines, making extinction of one or both species likely. High intrinsic growth rates result in higher reproductive rates when populations are below carrying capacity, reducing the probability of extinction and increasing the probability of stable coexistence at moderate levels of assortative mating and hybrid fitness. Very strong but incomplete assortative mating can induce low hybrid fitness via a mating disadvantage to rare genotypes, and this can stabilize coexistence of two species at high but incomplete levels of assortative mating. Given these results and evidence that it may take many millions of years of divergence before related species become sympatric, we postulate that coexistence of closely-related species is more often limited by insufficient assortative mating than by insufficient ecological differentiation.

## Introduction

Why do closely related species so often fail to co-occur? The question of coexistence of species is central to ecological and evolutionary sciences, although it is usually approached in different ways in the two fields (Germain et al. 2020). Ecologists have produced a rich body of work, referred to as niche theory or coexistence theory, to explain the conditions under which two species can be maintained within specific geographic areas (Vandermeer 1972; Chesson 2000; Siepielski and McPeek 2010; HilleRisLambers et al. 2012; Mittelbach and McGill 2019). Evolutionary biologists have approached this question through the lens of speciation theory (Panhuis et al. 2001; Turelli et al. 2001; Price 2008; Schluter and Pennell 2017), which examines the conditions that promote speciation, and via cline theory (Haldane 1948; Barton and Hewitt 1989; Polechová and Barton 2011; Gompert et al. 2017), which examines the dynamics of geographically structured hybrid zones between differentiated populations. While each of these approaches has generated profound insight into the causes of diversity, there is presently little integration due to their different assumptions regarding the amount of reproductive isolation (meaning less interbreeding than predicted by random mating, and/or low hybrid fitness). Coexistence theory generally assumes that the species in question are completely reproductively isolated. Speciation theory usually begins with a single species without any reproductive isolation and examines the conditions that cause the evolution of reproductive isolation (many speciation models do not end with complete reproductive isolation, instead producing a stable situation of strong but incomplete isolation (Servedio and Hermisson 2020)). Cline theory also assumes incomplete reproductive isolation, because it was developed to understand hybrid zones. Interbreeding (cross-mating) and hybridization (the production of hybrids) between populations that are otherwise fully recognized as distinct species is common (Barton and Hewitt 1989; Mallet 2005; Taylor and Larson 2019), suggesting that the potential for hybridization should be incorporated into species coexistence theory. Here, we ask what conditions are necessary to maintain two differentiated populations together in sympatry when there is incomplete reproductive isolation.

We envision what appears to be a common situation in nature: one species has been divided into two geographic regions where they have evolved some differences, and then these two populations have expanded their ranges into contact. They can differ genetically, ecologically, and in terms of mate preference. Hybrids might have lower fitness because of genetic incompatibilities or other disadvantages from having intermediate, mismatched, or transgressive ecological and behavioral traits. Here we present and analyze a model that incorporates such factors, asking under what conditions the two populations can coexist as distinct species. We focus on the simplest geographical situation: two differentiated populations come into full sympatry, without any spatial structure. We note that our model is not intended to precisely simulate specific situations seen in nature, which are invariably more complex than any simulation can be. Rather, our purpose is to show the coexistence outcomes that emerge from various combinations of assumed processes that are clearly specified. This approach results in much insight regarding the conditions necessary for coexistence.

For purposes of this analysis, if two differentiated populations maintain their distinctiveness in complete sympatry, we refer to them as distinct “species.” This is consistent with how taxonomy is usually practiced, even when there is some amount of hybridization and introgression. There are also many cases in nature of hybrid zones between otherwise geographically segregated taxa that are classified as different species, in which case the hybrid zone can play a role in preventing full sympatry. One goal of our analysis is to provide insight into what combinations of assortative mating and low hybrid fitness enable full sympatric coexistence of taxa classified as distinct species, rather than mere geographic segregration with a hybrid zone.

Current coexistence theory predicts stable coexistence when two fully reproductively isolated species differ sufficiently in resource use (‘stabilizing differences’) to counteract any differences in overall competitive ability (‘fitness differences’) (Chesson 2000; Germain et al. 2016; Mittelbach and McGill 2019). In our model, there are no differences in overall competitive ability between the populations, such that stable coexistence is expected when the populations are reproductively isolated and use entirely different resources. We note that our use of the words “stable” and “stability” are meant in an ecological sense rather than a mathematical sense: as stochastic simulations of finite populations would end in extinction given an infinite amount of time; we mean stable in the sense of a system tending to maintain certain characteristics for a long period of time despite minor stochastic perturbations (a similar definition to that used by Chesson 2000; Mittelbach and McGill 2019).

Incorporating hybridization into this framework requires specifying the resource use of hybrids and the resulting effect on the fitness of hybrids. This could be done in a way that favors hybrids, for instance if they use both resources as well as the parental groups do. It could alternatively be done in a way that penalizes hybrids, such that their potential to acquire resources is lower than the parental groups. Since our primary goal is to isolate the effects of assortative mating and intrinsic incompatibilities on species coexistence, we choose a method that does not give hybrids an advantage or disadvantage via resource use: individual ability to use each of two resources varies linearly with their genetic background, such that the total ability to use resources is constant among individuals (see Methods for details).

We use our model to address several key hypotheses regarding how hybridization may change the expectation of stable coexistence. First, we test whether a small amount of interbreeding (compared to none) can disrupt stable coexistence of two populations. Second, we test whether interbreeding and low hybrid fitness (compared to the fitness of the two starting populations) can result in extinction of one or both populations within the area of sympatry. Third, we test whether strong assortative mating can induce low fitness of hybrids, through rare-mating-type disadvantage. The results of our analysis lead us to conclude that interbreeding, hybridization, and/or reproductive interference (cross-population mating behaviour, without successful offspring production) likely play the major role in limiting sympatric coexistence between closely related species. We suggest that this largely explains a commonly-observed pattern in nature whereby sympatric coexistence of related species occurs only after a long period of allopatric differentiation, during which premating reproductive isolation must evolve to near completion (Price 2008; Weir and Price 2011).

## Methods

Our model assumes that two distinct populations have evolved elsewhere and come together into a single region with no spatial structure. We note that there could be other allopatric regions of each population, but the model assumes there is no gene flow with those regions and does not address them (see Irwin [2020] for a related model that does include such regions). The two starting populations have fixed genetic differences at a number of loci, and these genetic differences can be specified as determining differences in ecology, mating traits, and mating preferences, and also determining the fitness of hybrids. Our model is based on the Hybrid Zone with Assortative Mating (HZAM) model (Irwin 2020), which was designed to examine the role of assortative mating and low hybrid fitness in maintaining a narrow hybrid zone. We have modified this model in important ways, including removing spatial structure, adding ecological differentiation between the two initial populations, and tracking realized fitness (i.e., the average number of surviving offspring) of each genotypic group over time. The present model is designed to be able to separately examine effects of ecological differentiation and low hybrid fitness; this was done by ensuring no overall ecological advantage or disadvantage of intermediates due to ecological differentiation of the two initial populations (see below). There are two implementations of our model, one written in R (R Core Team 2021) and the other entirely re-written in Julia (Bezanson et al. 2017); the latter is faster by 1-2 orders of magnitude but does not yet have some of the options such as tracking realized fitnesses (see below). Careful comparisons were made to ensure the two implementations produced equivalent results (to see the R results graphs, see https://doi.org/10.1101/2021.04.04.438369 and compare those outcomes to those in the present paper, which were based on the Julia implementation except where noted).

This new model, named HZAM-Sym, is an individual-based simulation of two starting populations (A and B) in which all individuals have an equal probability of encountering all other individuals in the combined population. Individuals are diploid females or males (with equal numbers at the start of the simulation). One or more loci are assumed (in most cases that we present, there are 3), each with two alleles (designated 0 and 1, which are also their allelic values) that follow rules of Mendelian inheritance and are not physically linked nor sex linked. There is no mutation. At the beginning of each simulation, all population A individuals are 0/0 homozygotes and all population B individuals are 1/1 homozygotes, at every locus. These loci can be designated as affecting mating traits and preferences, ecological specialization (i.e., ability to use two resources), and the fitness of hybrids due to combinations of alleles from the two populations (see below for details). Loci that can affect these processes are called “functional loci” (*L* is the number of functional loci). Together, these loci produce a “functional trait” (*T*, ranging from zero to one) in an additive way, both between and within loci: to calculate *T*, the sum of all allelic values at functional loci is divided by the total number of alleles at functional loci (2*L*). In most simulations presented, a single trait (*T*) influences all mating traits and preferences, ability to use two resources, and the survival fitness of hybrids; however we also present some simulations where different sets of loci control different functional traits (see below).

Each simulation proceeds with cycles of mating, reproduction, and survival to adulthood, with non-overlapping generations. Distinct kinds of selection are incorporated into each step. These include mate choice (producing a pattern of assortative mating), density-dependent population regulation based on two resources (incorporated into the number of offspring of females, influenced by their functional trait), and differential survival probability to adulthood (with hybrid genotypes tending to have lower survival probability). We explain the rules of each in turn below.

### Mating

In the simplest case, mating is random, with each female being paired with a randomly chosen male. In the more interesting case, assortative mating is modeled through female choice based on the functional trait. This trait can be envisioned to be a male display and a female preference, but results are likely to be similar if those sex roles are reversed or if the trait is related to timing or breeding microhabitat rather than active choice.

Each female is presented with a random male who she can either accept as a mate or reject, in which case she is presented with another random male and repeats the process (excluding the previously rejected ones) until she accepts a male. Each female pairs with only a single male, regardless of her number of offspring (see below). Acceptance probability is determined by a comparison of female and male phenotypic trait values. If they are identical (*T*_diff_ = *T*_female_ – *T*_male_ = 0, where *T*_female_ is the trait of the female and *T*_male_ is the trait of the male), then she always accepts the male. If they differ, then probability of acceptance declines as their difference (|*T*_diff_|) increases, according to a Gaussian function with standard deviation *σ*_pref_ (fig. 1A; for a case where empirical mating patterns based on size are similar to this Gaussian function, see Perini et al. 2020). This female choice system means that there is variation among males in their number of mates (with some having no mates) whereas almost every female has one mate (the one very rare exception being when a female rejects all males currently in the simulation, in which case she does not produce any offspring—this happens only when one of the founding populations is near extinction). The strength of assortative mating (*S*_AM_), which is directly related to σ_pref_, is expressed as the ratio of the probability of a female accepting a presented male that is identical to her (*T*_diff_ = 0) to the probability of a female accepting a presented male that is one unit of phenotype different from her (i.e., a full heterospecific, *T*_diff_ = 1). Hence if *S*_AM_ = 1, then there is no difference in probability of acceptance; if *S*_AM_ is 1000, then a female has a 1000 times greater probability of acceptance of an identical male compared to a fully different male.

**Figure 1.**
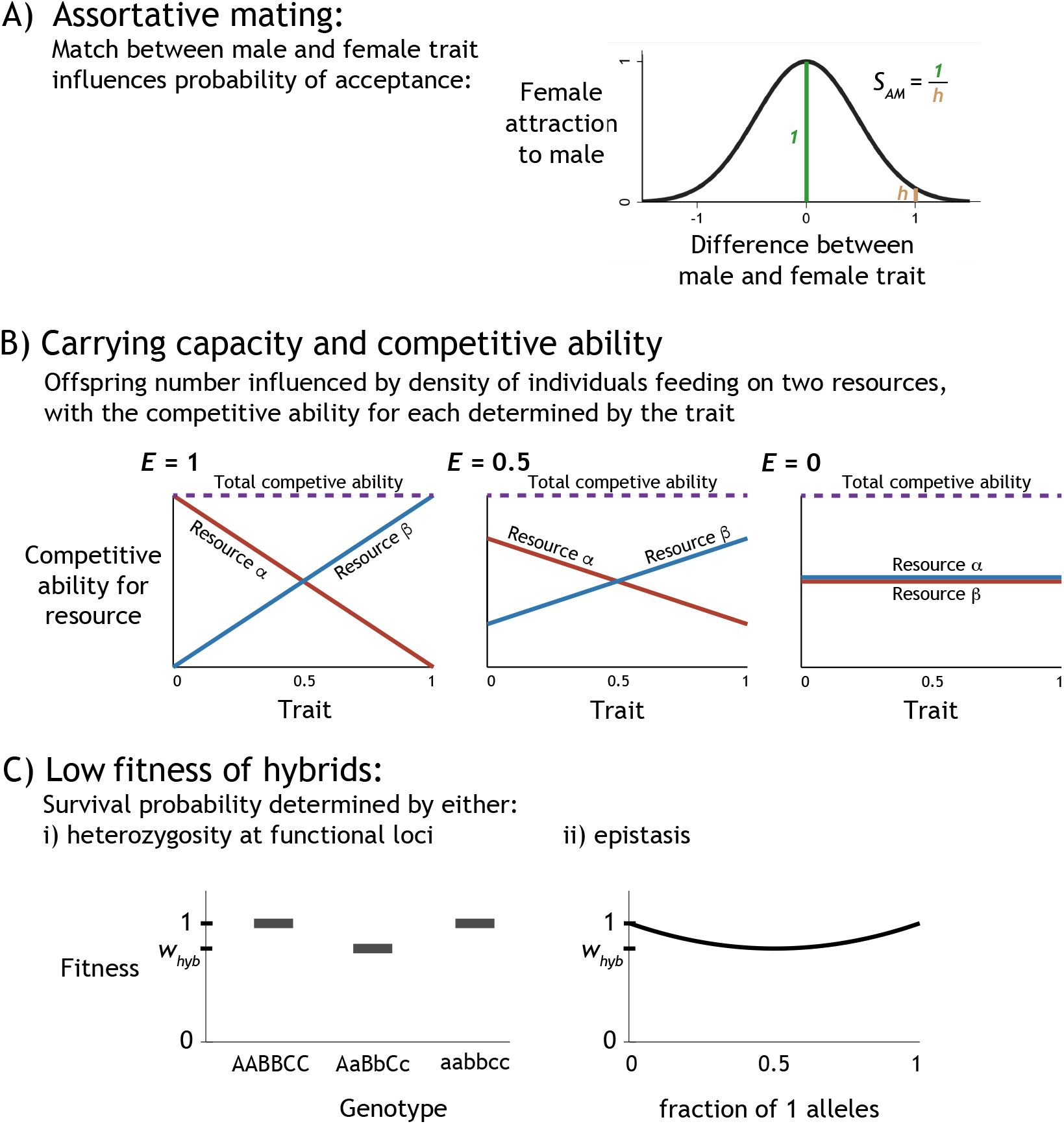
Three effects of functional loci that together determine a functional trait: (A) the degree of mismatch between a female and candidate male mate determines the probability that she will reject him and encounter a different male instead. (B) The functional trait determines competitive ability of the individual for two resources, which influences the expected number of offspring produced by each female. (C) Reduced probability of survival to breeding age can depend on i) heterozygosity at functional loci or 2) epistasis, including interactions both between and within loci.

### Reproduction based on density dependence via two resources

Ecological differentiation is modeled as trait-dependent variation in the competitive ability to use two resources (α and β, with fixed carrying capacities *K_α_* and *K_β_* respectively). We refer to these competitive abilities as *U_α_* and *U_β_*, which are both characteristics of each individual. In the simplest case of full ecological differentiation, we specified that individuals of starting population A (i.e., *T* = 0) can use resource α but not β, such that *U*_α_ = 1 and *U*_β_ = 0. Likewise, individuals of starting population B (i.e., *T* = 1) can use resource β but not α, such that *U*_α_ = 0 and *U*_β_ = 1. Intermediate individuals (i.e., produced through hybridization) have an intermediate competitive ability on each. As the trait value increases from 0 to 1, the competitive ability for resource α declines linearly and the competitive ability for resource β increases linearly (fig. 1B, left panel).

This method of modeling ecological differentiation results in the total competitive ability (the sum of competitive abilities on resource α and resource β) being the same (equal to one) for all trait values (fig. 1B). This approach avoids an ecological disadvantage or advantage of intermediate forms. There is still a diversity-promoting role for ecology, manifested in frequency-dependence: If most individuals have trait 0, then competition for resource α is greater than competition for resource β, causing individuals with trait 1 to have higher fitness than those of trait 0. This formulation does not favor or hinder bimodality or unimodality of the total population (i.e., sum of all individuals of both populations and hybrids). Any total population that has an average trait of 0.5 has the same resulting carrying capacity. For example, one hybrid population all with phenotype 0.5 has the same carrying capacity as a population consisting of 50% trait 0 and 50% trait 1.

The model can consider varying degrees of ecological differentiation, using a parameter *E* which can take values from 0 to 1. In the full differentiation described above, *E* = 1 (fig. 1B, left panel). When there is less ecological differentiation (*E* < 1; fig. 1B, middle and right panels), for population A the competitive ability on resource α (*U*_α_) is equal to 0.5 + *E*/2 and on resource β (*U*_β_) is 0.5 - *E*/2. Similarly, for population B the competitive ability on resource α (*U*_α_) is equal to 0.5 - *E*/2 and on resource β (*U*_β_) is 0.5 + *E*/2. Competitive abilities of intermediate trait values are determined by linear relationships between these values. In the case of no ecological differentiation (*E* = 0), these formulae result in all trait values having a competitive ability of 0.5 on both resources (fig. 1B, right panel). This mathematical approach encapsulates the idea that individuals who can utilize two resources are half as good at consuming a single resource compare to an individual that is specialized only on that one resource.

Before the reproduction phase of each generation, the sums of competitive abilities of all individuals are calculated, for each resource. We call these sums *N*_α_ and *N*_β_, for resources α and β respectively, as they correspond to the equivalent number of perfectly suited individuals using the resource. These can be thought of as representing the intensity of resource use by the entire population.

For each of the two resources (α and β) we then calculate expected population growth rates (*r*_α_ and *r*_β_) of the combined consumer population (i.e., both species and the hybrids), based on the carrying capacity of each resource (*K*_α_ and *K*_β_), the intrinsic growth rate of the consumer population when small (*R*), and the intensity of resource use (*N*_α_ or *N*_β_). We use the discrete time analog of the continuous logistic growth equation (Prout 1978; Liou and Price 1994). The growth rate of the consumer population due to each resource is given by:

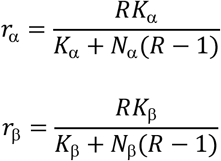

Each female’s expected number of offspring (*c*) is then a function of her ability to use the two resources (*U*_α_ and *U*_β_) and the consumer population growth rates due to each resource:

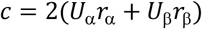

The 2 in this equation is because males do not directly produce offspring but are produced by mothers. After the expected offspring of each breeding female is calculated in this way, her actual number of offspring is determined by a random draw from a Poisson distribution with mean *c*.

At each genetic locus, offspring receive one allele from the mother and one from the father, each chosen randomly. Sex of each offspring is chosen randomly, with 50% probability of each sex.

### Survival

Low fitness of hybrids is modeled as reduced survival to adulthood (fig. 1C), based on either underdominance (i.e., heterozygote disadvantage), or epistasis (which includes both underdominance and between-locus incompatibilities). In both, complete heterozygotes at all functional loci (i.e., F1 hybrids) have survival probability *w*_hyb_, whereas complete homozygotes (i.e., members of the starting populations A and B) all survive to adulthood (i.e., survival probability equal to 1). In the underdominance-based fitness, the effects of different loci on survival fitness are assumed to be equal and multiplicative, such that for genotypes with only some heterozygous loci the probability of survival is:

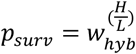

where H is the number of heterozygous functional loci and L is the total number of functional loci. In the epistasis-based fitness, the probability of survival is determined following Barton and Gale (1993):

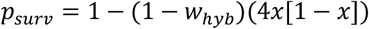

where x is the total fraction of 1 (or 0) alleles at all functional loci.

### Running simulations and categorizing outcomes

Most simulations presented here use three functional loci and begin with a total of 1000 individuals, in two populations each of 500 individuals (divided equally between males and females). We present results of simulations at all combinations of assortative mating *S*_AM_ = {1, 3, 10, 30, 100, 300, 1000, complete} and *w*_hyb_ = {0, 0.1, 0.2, 0.3, 0.4, 0.5, 0.6, 0.7, 0.8, 0.9, 0.95, 0.98, 1}, with 25 replicates per combination. These results are presented for various combinations of *E* = {0, 0.25, 0.5, 0.75, 1} and *R* = {1.025, 1.05, 1.2, 2.6}, for both underdominance and epistasis, and with other special cases as described below.

Each simulation is classified into one of four outcome types based on the state of the simulated total population after 1000 generations (or as otherwise specified). To do this, we first calculate each individual’s proportion of alleles that are allele 1, which we refer to as the hybrid index (HI) of that individual. We then calculate the proportion of final individuals that have more than 90% of their alleles from population A (that is, HI < 0.1) and the proportion that have more than 90% of their alleles from population B (that is, HI > 0.9). We then categorize outcomes according to these criteria:

- Extinction, if no individuals (of both starting populations and hybrids) are present at the end of the simulation.
- One species, if at least 85% of the individuals have HI < 0.1 or if at least 85% have HI > 0.9.
- Two species, if more than 15% of individuals have HI < 0.1 and more than 15% have HI > 0.9 and the sum of those having HI < 0.1 or HI > 0.9 is more than 85% of the population.
- A blended population, in all other cases.

### Confirmation of necessity of ecological differentiation for stable coexistence

Initial testing of the model’s behavior was done to determine an appropriate population size and run length (i.e., number of generations) that would suitably demonstrate what parameter combinations lead to long-term coexistence. These were done for the case when the two species are ecologically identical (*E* = 0) and the case when the species are completely differentiated on different resources (*E* = 1). We decided that a starting population of 1000 individuals (*K*_α_ = *K*_β_ = 500) and a generation time of 1000 is generally sufficient to distinguish long-term coexistence from other outcomes.

When there is no ecological differentiation (*E* = 0) and complete assortative mating (no hybridization), the two initial populations persist for some time but eventually one or the other goes extinct (fig. 2A). This is a result of them being ecologically identical and finite in population size, such that they are jointly regulated by a single carrying capacity. Chance variations in their population sizes eventually lead to one going extinct. This phenomenon is well understood, often referred to as “unstable coexistence” (e.g., (Chesson 2000; Mittelbach and McGill 2019)) but perhaps better referred to as transient co-occurrence due to neutrality (Germain et al. 2020)).

**Figure 2.**
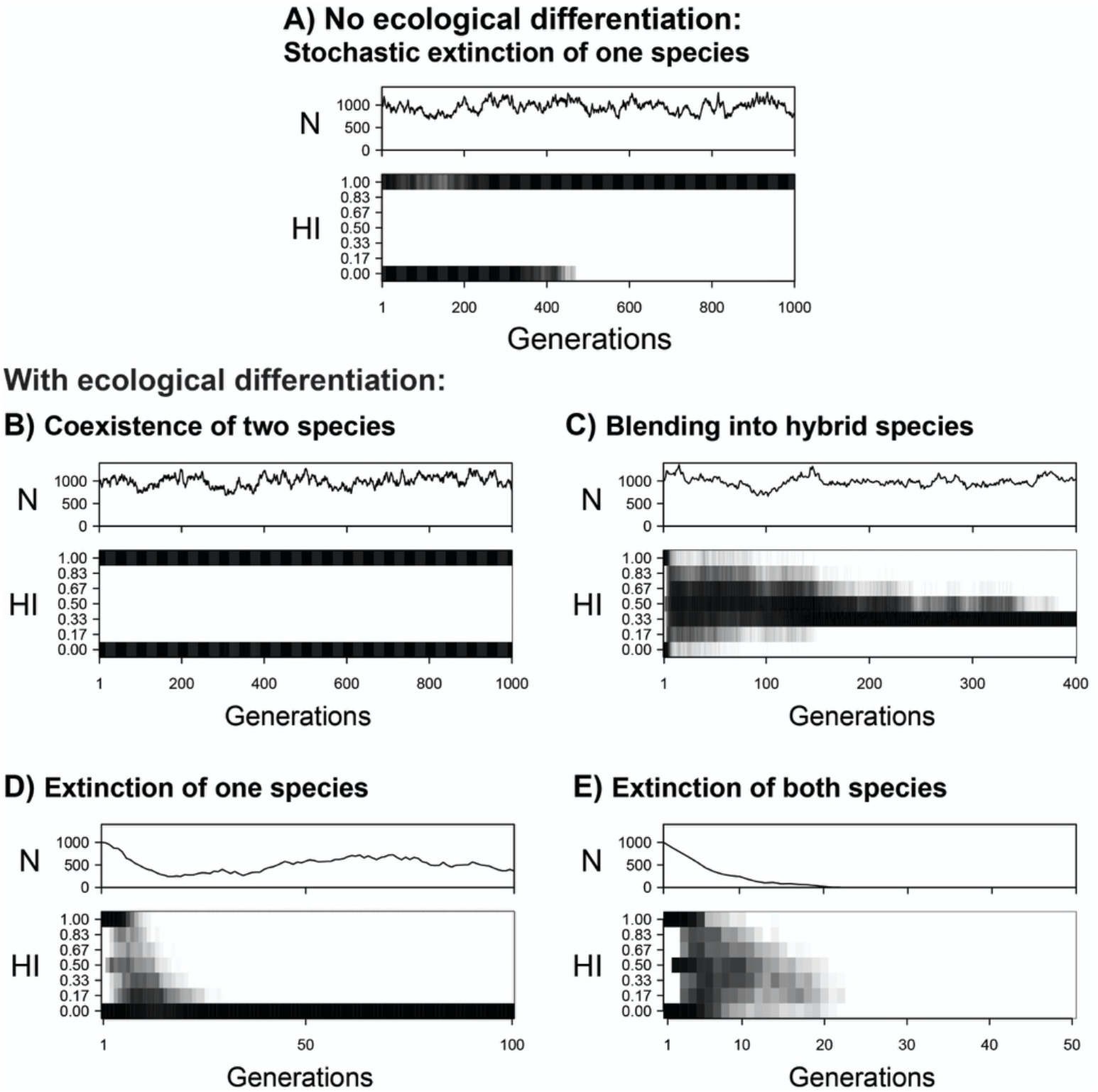
Five example simulations illustrating various outcomes of contact between two species. In each case, three Mendelian loci additively determine a genetic and phenotypic hybrid index (HI) ranging from 0 (one of the initial species) to 1 (the other initial species). For each simulation, the population size (*N*) and the density of each HI category is shown for each generation. In (A), no ecological differentiation (*E* = 0) and complete assortative mating result in stochastic loss of one species. In (B-E), there is ecological differentiation (*E* = 1) of the two species. In (B), the ecological differentiation enables long-term coexistence of two species that have complete assortative mating. In (C), when hybrids have the same fitness as the starting populations (*w*_hyb_ = 1), assortative mating of 10x (*S*_AM_ = 10) is insufficient to prevent collapse of the two species into a hybrid form. The same settings are used in (D), except hybrid fitness (*w*_hyb_) is lowered from 1 to 0.6— this leads to population decline following hybridization and selection leading to recovery of just a single original species. The same settings are used in (E), except assortative mating is reduced to 3x—these conditions lead to complete extinction of both species and hybrids. The R language implementation of HZAM-Sym was used for all simulations in this figure, with *R* = 1.05, *K*_α_ = 500, *K*_β_ = 500, and the underdominance method of specifying survival fitness.

With ecological differentiation, long-term stable coexistence is possible. In the case of complete specialization on different resources (*E* = 1) and complete assortative mating, the two populations (species, in this case) persist for the long term because they are regulated by carrying capacities for two different resources (fig. 2B).

## Results

The simulations reveal that even a small rate of interbreeding can dramatically alter the conditions under which two differentiated populations can persist in sympatry. For example, if we start with the conditions modelled in figure 2B (complete ecological differentiation) and reduce the strength of assortative mating from complete to merely strong (10x assortative mating; *S*_AM_ = 10; meaning 10x stronger preference for conspecific over heterospecific), blending into a single hybrid species (with intermediate genotypes) occurs within the first 10 generations after contact (fig. 2C). Although formation of hybrids is initially rare, assortative mating means that hybrids tend to mate with other hybrids (if they are sufficiently common to encounter each other). This, combined with the fact that the initial populations tend occasionally to mate with each other or with hybrids, means that intermediates tend to build up over time and the initial extreme genotypes decline. Extremely high levels of assortative mating are needed to maintain two species. In the case of *R* = 1.05, roughly 300x assortative mating is required (fig. 3B).

**Figure 3.**
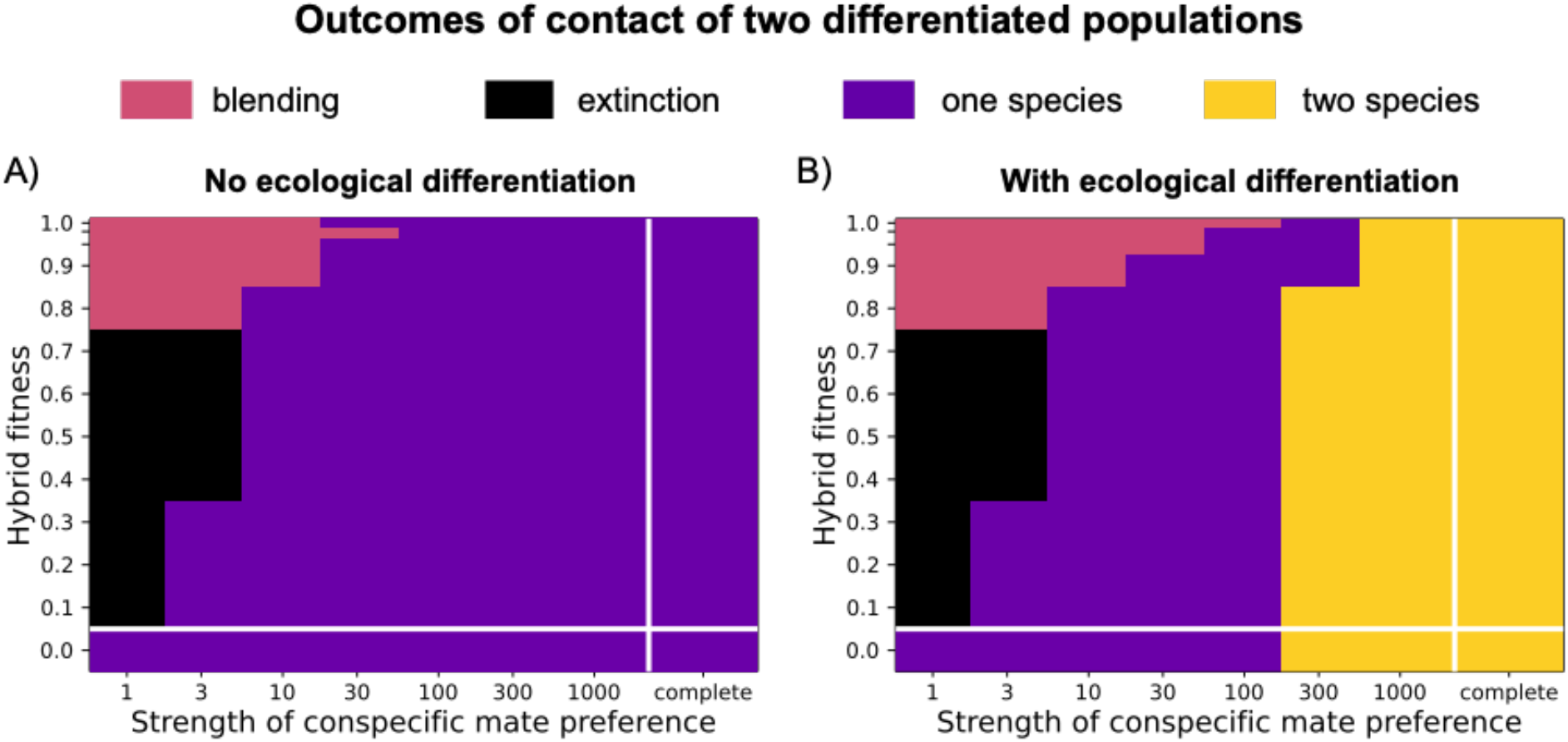
Outcomes of contact between two populations after 1000 generations (A) without ecological differentiation and (B) with full ecological differentiation, at various combinations of assortative mating strength (along horizontal axis) and hybrid fitness (vertical axis). Twenty-five simulations were run (using the Julia language implementation) under each set of parameter combinations (as indicated by the marks along each axis), for a total of 5200 simulations for this figure. Colors represent the most common outcome for each set: Black = extinction of both species; Purple = one species remaining (extinction of the other); Salmon = Hybrid population; Yellow = Two species; ties for most common outcome (rare with 25 simulations) were decided by a random draw from the most common. To see a detailed breakdown of the frequency of outcomes under each set, see figure S1. In these simulations total carrying capacity is 1000 (500 on each of two resources) and growth rate *R* = 1.05. When no ecological differentiation of the starting populations (*E* = 0, in A), one or the other of them is lost within 1000 generations. When there is full specialization on different resources (*E* = 1, in B), then strong assortative mating allows the maintenance of two species. These results are based on the heterozygote disadvantage method of modelling hybrid fitness; for results based on epistasis, see figures S2, S3.

Our second result is that hybridization with low hybrid fitness often leads to extinction of one or both populations. An example of extinction of one population is illustrated in figure 2D, which has the same conditions as figure 2C (10x assortative mating, *E* = 1, *R* = 1.05) except with hybrid fitness (*w*_hyb_) reduced from 1 to 0.6, working against the buildup of intermediate genotypes. There is then a tension between cross-mating that is producing some intermediates and selection that is favoring the extremes. The tension is resolved by the distribution moving toward one of the original extreme genotypes, recovering one of the parental populations but causing extinction of the other. This happens because when one of the parental populations happens to become rarer than the other, a higher fraction of the rarer population is producing low-fitness hybrids than the more common population, hastening the demise of the rare population. In the top panel of figure 2D we see how the population size (*N*) begins at a combined carrying capacity of 1000 (500 for each parental species) but declines due to the low fitness of hybrids and then rises as a single high-fitness parental population is recovered. After this one-population outcome is established, the population fluctuates around the single-population carrying capacity of 500. The overall system went from two differentiated populations specialized on two different resources to a single population specialized on one resource. At the end of the simulation, one resource is now not used at all, and the overall population size is half the value at the start.

When low fitness of hybrids causes an even more severe population decline, the result can be complete extinction of both populations (and all intermediates). An example is seen in figure 2E, which has the same parameter settings as figure 2D except with assortative mating reduced from 10x to 3x (*S*_AM_ = 3). This change causes low-fitness hybrids to be produced at a higher rate, and the decline is so severe that the combined population cannot evolve fast enough to avoid extinction.

Figure 3 illustrates how hybrid fitness, the strength of assortative mating, and ecological differentiation interact to influence species cooccurrence (colors are color-blind-friendly, based on the “plasma” color scale; Garnier 2018). We see that the case of no ecological differentiation (*E* = 0; fig. 3A) and the case of ecological differentiation (*E* = 1; fig. 3B) differ in that only the latter has some parameter space in which two species are likely to be present after 1000 generations. This however requires strong assortative mating. At more moderate levels of assortative mating, the outcome tends to be one remaining starting population. At lower levels of assortative mating, the outcome depends on hybrid fitness: high hybrid fitness tends to lead to blending, whereas lower hybrid fitness leads to extinction of both populations and their hybrids. Note that extinction of one or both populations is likely even when hybrids have zero fitness. In this case, reproductive isolation is complete and the two species are ecologically differentiated—but even so, very high assortative mating is required for long-term sympatric coexistence. This is because the production of zero-fitness hybrids consumes reproductive effort, resulting in population decline if the rate of cross-mating is sufficiently high.

A crucial parameter in influencing these outcomes is *R*, the intrinsic growth rate. In figure 4, we see that a higher intrinsic growth rate leads to less parameter space over which extinction of one or both populations occurs, and more parameter space over which blending or coexistence occurs. This can be understood as a result of a higher intrinsic growth rates resulting in less potential for low hybrid fitness to reduce population size. Higher population sizes provide selection with more time and power to influence the outcome.

**Figure 4.**
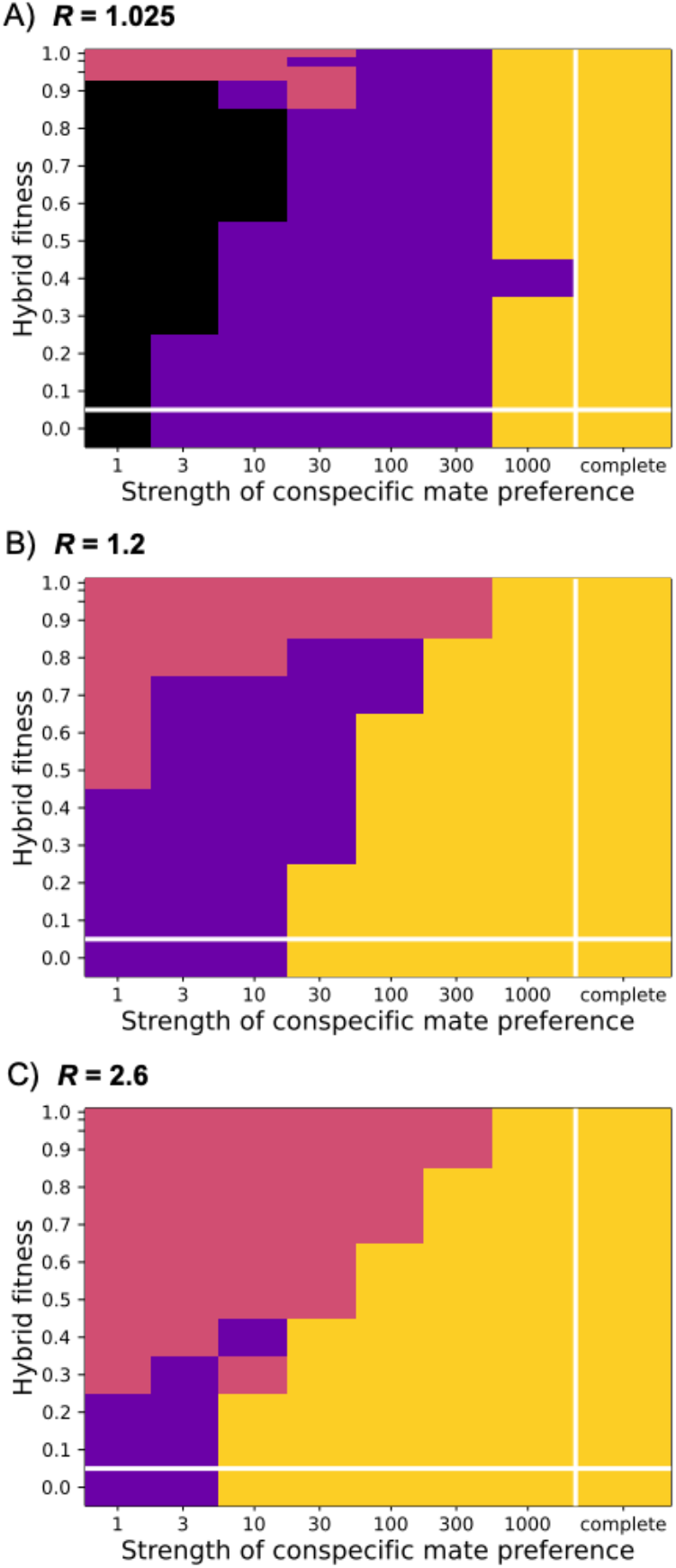
Higher intrinsic growth rate reduces the probability of extinction of one or both species, increasing the parameter space over which two species can be maintained. See the caption of figure 3 for full explanation of colors and figure format. Each panel was produced using full ecological differentiation (*E* = 1), total carrying capacity of 1000, and 1000 generations, and the underdominance method of hybrid fitness. Intrinsic growth rates (*R*) in each panel were (A) 1.025, (B) 1.2, and (C) 2.6. Twenty-five simulations were run under each set of parameter combinations (7800 simulations total in this figure), with colors representing the most frequent outcome for each set: salmon = hybrid population; black = extinction of both populations; purple = one population remaining (extinction of the other); yellow = two species. To see a detailed breakdown of the frequencies of outcomes, see figure S4. For results using the epistasis method of hybrid fitness, see figures S5, S6.

Our third main result is that very strong choice-based assortative mating (e.g., *S*_AM_ above 300 or so) can induce low fitness of hybrids due to their rarity as mating types, a type of sexual selection against hybrids, and this can influence whether the populations coexist. We tracked the average number of offspring produced per each trait class during each generation of the simulations. Figure 5A shows a simulation with no intrinsic differences in fitness between hybrids and initial populations (*w*_hyb_ = 1), strong assortative mating (*S*_AM_ = 1000), high ecological differentiation (*E* = 1), and *R* = 1.2. Hybrids are produced throughout the simulation, but their realized fitness is much lower than the initial populations (e.g., fitness is about 70% for F1 hybrids—see the line for HI = 0.5). This low fitness is due to males suffering a rare-mating-type disadvantage because of the rarity of females with similar trait values. Most females are of one of the initial genotypes, and they strongly prefer males of their own type. Under these conditions, two species coexist long-term. In figure 5B, we see a simulation under similar conditions except the strength of assortative mating is reduced to *S*_AM_ = 300. For the first hundred generations or so, hybrids are produced at a low rate but tend to have low fitness. Rare hybridization and backcrossing however causes some gene exchange between populations, such that partially intermediate trait classes (e.g., *T* = {0.17, 0.83}) gradually rise in frequency. Eventually, there is enough of a continuum of types that hybrid fitness rises—hybrid males encounter more females with similar trait values. There is a transition to a different fitness landscape, with intermediates now having higher fitness than the extremes of the trait distribution. This eventually leads to loss of variation, with the system converging on a single hybrid population with no genetic variation. In this case, strong assortative mating has a different impact at different stages of the simulation: when there are two discrete populations, strong assortative mating tends to cause low fitness of hybrids; when hybrids are more common, the same assortative mating tends to cause higher fitness of hybrids and eliminate the extreme (initial) phenotypes.

**Figure 5.**
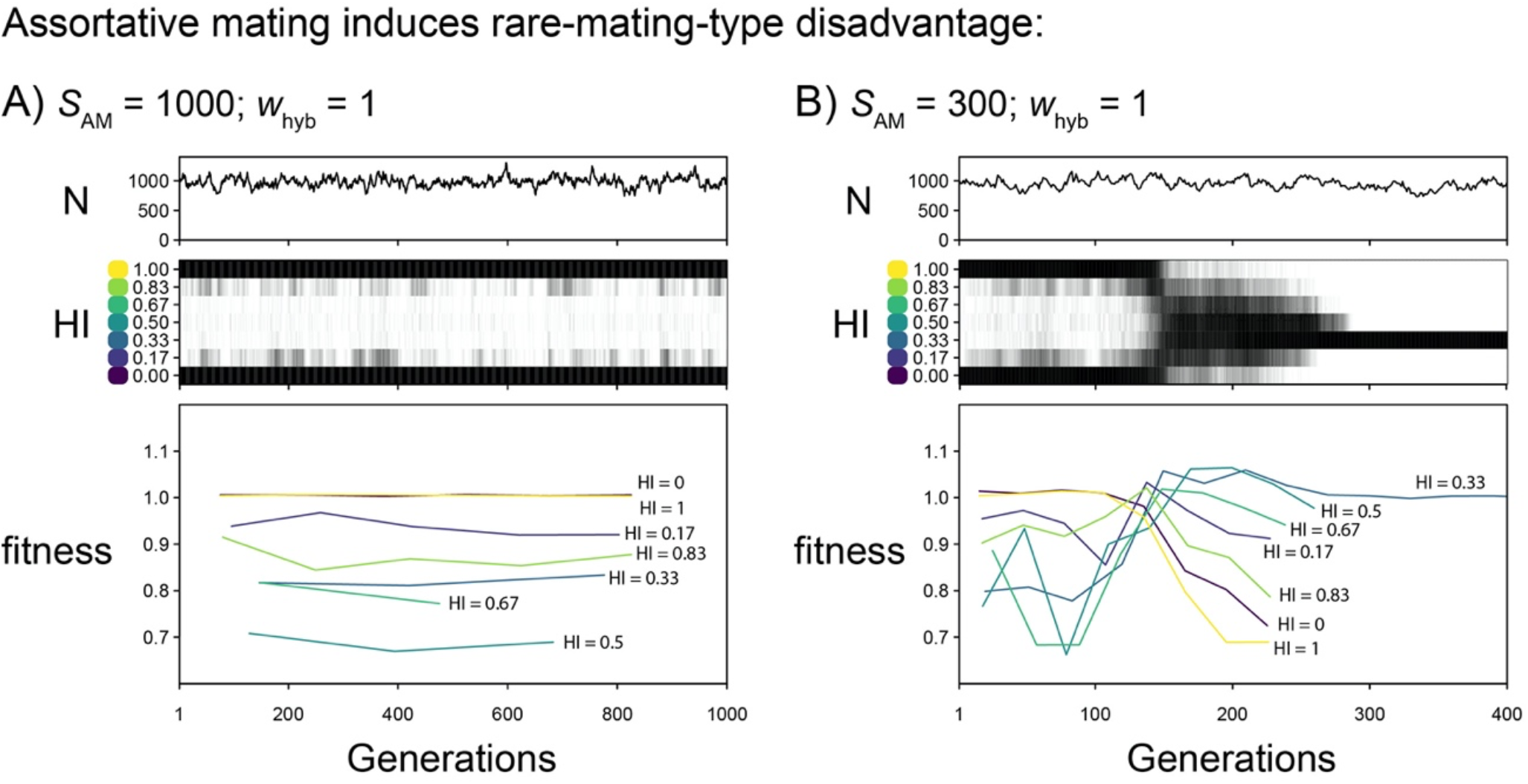
Assortative mating can cause lower fitness in rare mating types, favoring common mating types. In these two example simulations, there is ecological differentiation (*E* = 1), intrinsic growth rate *R* = 1.2, and no pre-determined lower fitness of hybrids (*w*_hyb_ = 1). The only difference is in the strength of assortative mating: 1000x in A) and 300x in B). In A), there is some rare hybridization and backcrossing throughout the simulation, but the fitness of hybrids is low (i.e., hybrids produce fewer offspring on average) due to low average attractiveness of rare males as potential mates—they rarely encounter females with similar trait values. In B), for the first 100 generations or so, hybrids have low fitness compared to the parental species, keeping hybrid classes rare. At about generation 150, intermediate classes become common enough that they no longer suffer a rare-mating-type disadvantage, and the fitnesses then flip, with intermediate classes being the common types and the extreme parental traits having low fitness. A hybrid species with no variation emerges. Produced using the R language implementation of HZAM-Sym.

Because of the way that assortative mating based on mate choice can cause frequency-dependent selection against rare mating types, some mathematical models of assortative mating have removed this sexual selection by using group-based mating (Felsenstein 1981; Otto et al. 2008) or an approach that is designed to neutralize the sexual selection (De Cara et al. 2008; Pennings et al. 2008). To explore whether our main results hold when there is assortative mating without sexual selection, we changed the choice-based mating system into a group-based mating system where individuals join groups based on their functional traits and there is random mating within the groups. Imperfect assortment is modeled through movement between the groups prior to mating. Full methods and results of this approach are described in the Appendix. Under this group-based mating, the first two results that we have reported are even more strongly supported: two species cannot coexist even under the lowest non-zero rate of hybridization tested, and extinction of both species occurs over an even broader set of conditions than in the choice-based mating system (fig. S1). We think this group-based mating system is less realistic than the choice-based system, hence we emphasize the choice-based system in our analysis.

To examine how robust our conclusions are to the modelling decisions used, we explore how deviating from those affected the results. Our main simulations use 1000 generations, whereas figure S7 shows the results of runs using different numbers of generations, ranging from 125 to 4000. Between 125 and 500 generations there is some modest decline in the parameter space where two species coexist, but beyond that such change is minor (fig. S7).

To specify low survival fitness, we use heterozygote disadvantage in the main simulations. Using a model of epistasis, which also includes between-locus interactions, results in a similar boundary between the parameter space where two species coexist vs. other outcomes (figs. S2, S5; compare to figs. 3, 4). However, there is a reduction in the prevalence of the blending outcome vs. the one-species outcome. This is a result of the heterozygote disadvantage model having the possibility of recovering high fitness through establishing homozygosity for different starting populations at different alleles.

Our main simulations used 3 loci, whereas figure S8 shows results for numbers of loci ranging from 1 to 27. There is a sizeable effect of the number of loci, with more loci causing a smaller region of parameter space where two species coexist and more parameter space where full extinction occurs. This result is likely to be an effect of the strength of selection on each locus: for a given total strength of selection against hybrids (1 – w_hyb_), more loci mean weaker selection on each locus, lowering the effectiveness of selection in countering the buildup of intermediates.

In the main simulations, each of the loci jointly contribute equally to female preferences, male signaling, low hybrid fitness, and resource competition, analogous to real situations where loci encode a single trait that has multiple effects (e.g., in *Littorina*; Perini et al. 2020). We also conducted simulations where two separate sets of loci (3 loci each) controlled female and male traits (fig. S9A), and others where three sets of loci (3 loci each) controlled female traits, male traits, and ecological traits (meaning the low hybrid fitness and the resource competition trait; fig. S9B). Both resulted in less parameter space where long-term coexistence of two species occurs, and more where full extinction occurs (compare fig. S9 with fig. 3B). This is because the different sets of loci can become uncoupled through reproduction of hybrids, making selection less effecting at maintaining two differentiated forms.

We also explored the effects of unequal initial population sizes (rather than the 1:1 ratio in the simulations described above). Figure S10 shows that there is an effect of imbalanced starting ratios on the boundaries in parameter space between different outcomes. The two-species outcome covers slightly less parameter space than in the 1:1 starting case, because an initially rare population tends to be more subject to rare-mating-type disadvantage, the cost of production of low-fitness of hybrids, and stochastic loss. The blended and extinction outcomes also cover less parameter space, because the hybrid population becomes more likely to converge quickly on one of the starting genotypes (the one-species outcome) when allele frequencies are highly imbalanced.

## Discussion

These results indicate that interbreeding can greatly limit the ability of species to coexist in the same geographic area, while also revealing parameter combinations where maintenance of two sympatric species is possible despite incomplete reproductive isolation. When two species use entirely different environmental resources but have no differences in intrinsic growth rates or sensitivities to other factors, current coexistence theory predicts stable coexistence (Chesson 2000; Mittelbach and McGill 2019). These are the conditions modelled in the bulk of the simulations presented here (*E* = 1 in all figures except in fig. 3A and parts of fig. 6). Yet when we allow assortative mating to be incomplete, outcomes other than coexistence are observed under a wide range of parameter space. These other outcomes include blending into a single hybrid species or hybridization-induced extinction of one or both initial populations. These other possibilities start to become likely when assortative mating is reduced from complete to merely extremely strong, for example a 300 times greater preference (when *w*_hyb_ = 1) of a female for a male of her own species compared to a male of the other species. Collapse of two populations into a hybrid population despite strong assortative mating has also been observed using other modelling approaches (Singhal and Moritz 2012; Pulido-Santacruz et al. 2018; Cronemberger et al. 2020; Irwin 2020), although the approach used here differs from earlier approaches by incorporating ecological differentiation and the possibility of population decline.

**Figure 6.**
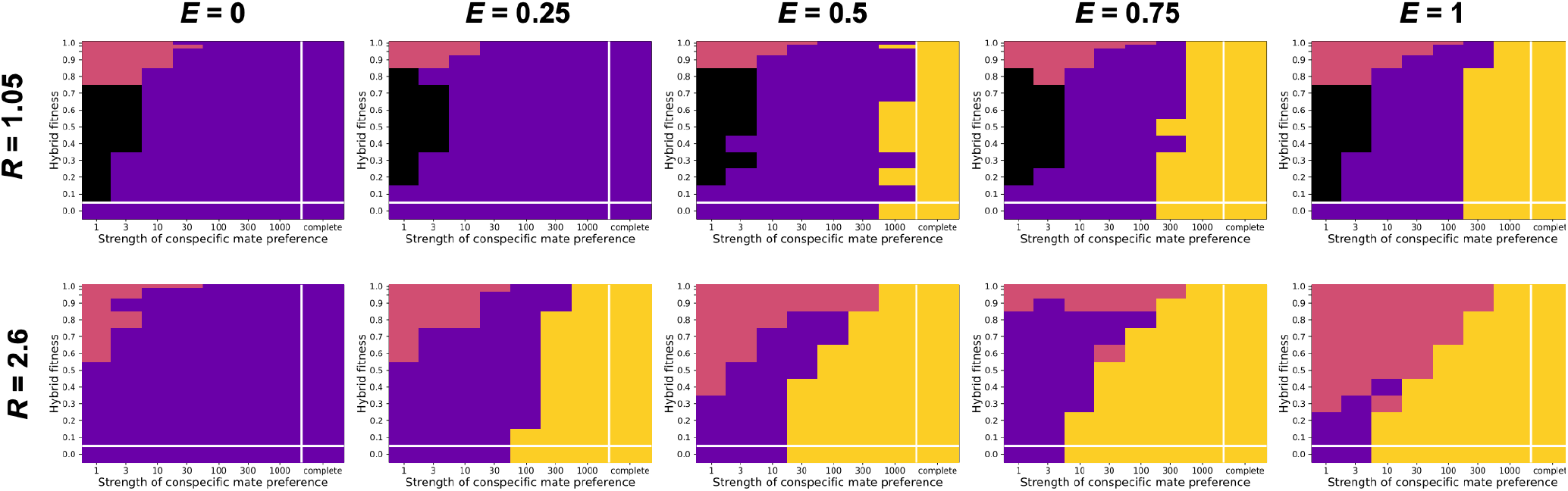
Coexistence of two species (yellow) versus other outcomes (black = extinction; salmon = blending; purple = one species) is jointly determined by the intrinsic growth rate (*R*), the amount of ecological differentiation (*E*), the strength of conspecific mate preference, and hybrid fitness. Each panel has the same format (e.g., axis labels) as figures 3 and 4, using the *R* shown left of each row and the *E* shown above each column. Twenty-five simulations were run under each set of parameter combinations, using the Julia implementation and the underdominance method of survival probability, with colors representing the most frequent outcome for each set.

The boundary conditions for stable coexistence depends not just on the strength of assortative mating, but also on hybrid fitness (*w*_hyb_) and intrinsic growth rate (*R*). As hybrid fitness is decreased, the boundary between stable coexistence and other outcomes is observed at decreasing assortative mating strengths (e.g., fig. 4B). This can be understood as a result of the low hybrid fitness working against the buildup of intermediate types, allowing coexistence of two species at moderately high levels of assortative mating. Still, under all the conditions modelled here, maintenance of two species is not the likely outcome when assortative mating is less than 10x, an amount that would be considered quite strong compared to that observed in many studies of hybrid zones (Irwin 2020).

A particularly interesting scenario is when hybrids between two populations have zero fitness, such that the two populations are by definition completely reproductively isolated (i.e., no possibility of genetic exchange) and thereby considered distinct species. These simulations illustrate a way in which two such species may be unable to coexist even if ecologically divergent: if assortative mating is not strong (e.g., less than 100x in the case of *E* = 1 and *R* = 1.05; fig. 3B), then reproductive interference (i.e., the cross-mating of two species, requiring reproductive effort and possibly the production of zero-fitness hybrids; Kuno 1992; Whitton et al. 2017) leads to population decline and extinction of one or both species. Hence species coexistence is not assured despite complete ecological differentiation and complete reproductive isolation. A possible example is the Pacific Wren and Winter Wren species pair, which interbreed and produce F1 hybrids that have zero fitness (Mikkelsen and Irwin 2021). Their geographic ranges come into close contact but with little overlap and low population density where they co-occur, consistent with reproductive interference leading to a failure to coexist (Mikkelsen and Irwin 2021).

The role of intrinsic growth rate (*R*) in influencing the outcome of contact between two hybridizing populations has received little previous attention. This is largely because much hybrid zone theory is based on mathematical models that assume constant population sizes and/or densities (Durrett et al. 2000; Polechová and Barton 2011), in some cases infinite (Haldane 1948; Barton and Hewitt 1989). In those cases, fitnesses are relative and the population is not allowed to decline in size. In the HZAM model presented here, survival fitnesses are absolute rather than relative, such that low average survival fitness can result in population decline. Working against this population decline is the density-dependent increase in reproduction when populations are below their carrying capacity. The intrinsic growth rate determines the magnitude of this increase in reproductive rate: a higher *R* results in reduced parameter space over which extinction of one or both populations is the likely outcome. Species coexistence then becomes possible over more combinations of lower hybrid fitness and reduced strength of assortative mating (i.e., the lower-middle parts of the panels in fig. 4).

The ultimate outcome of continual population decline is extinction, which is observed for one or both species over much of the explored parameter space. When intrinsic growth is low and assortative mating strength is low, extinction of both populations (and all hybrids) can occur over a wide range of low hybrid fitnesses (fig. 4A). Increasing assortative mating to moderate levels results in a higher probability of one population persisting—this is because assortative mating slows the rate of production of low-fitness hybrids and thereby reduces the rate of population decline, giving time for selection to shape the combined population back to a single starting genotype (compare figs. 3D and 3E).

One factor that is not incorporated into the present analysis is search cost. If individuals pay a cost per each potential mate they reject or each time they are rejected (via time and/or energy expended in the interaction), this could have complex effects on the coexistence outcomes. Search costs are expected to impact rare mating types more than common types, such that they may in some cases promote coexistence of two species by reducing fitness of hybrids. However, they are also expected to reduce fitness of the overall population and especially either starting genotype that happens to become more rare, leading to greater instability of the system. Adding search cost to these models would be worthy in future development of these models, but will require much thought as it is not clear that per-reject costs should be the same for all levels of assortative mating strength.

Our results help explain a common pattern in nature: closely related species tend to have geographic ranges that are either allopatric (no geographic overlap) or parapatric (separate ranges that are in contact), whereas true sympatry (broad overlap of ranges) typically occurs only after millions of years of divergence (Weir and Price 2011). Explanations for this long span of time before sympatry include long-term geographic barriers, competitive exclusion due to similar niches, and lack of reproductive isolation (Weir and Price 2011). In the models presented here, there is no geographic barrier and no niche overlap between the species (when *E* = 1), revealing the power of incomplete reproductive isolation alone to limit sympatric coexistence. This power is remarkable: lowering the strength of assortative mating from complete to merely 300 times greater preference for conspecifics (compared to heterospecifics) results in collapse of the two populations into either a hybrid population or just one of the original species. If low hybrid fitness is also a factor, then extinction of both populations becomes a possibility (at lower strengths of assortative mating). This possibility of extinction of both populations, or extinction of one with the remaining one at half the total carrying capacity, may help explain an often-observed pattern of gaps in distribution or areas of low density within spatially structured hybrid zones (Mikkelsen and Irwin 2021).

Assortative mating has usually been thought of as a powerful “prezygotic barrier” to gene flow, but our results show its effect on coexistence can be dependent on its role as a “postzygotic barrier” lowering hybrid fitness due to rare-mating-type disadvantage. When hybrids have survival fitness equivalent to the starting genotypes (*w*_hyb_ = 1), assortative mating must be extremely strong (e.g., *S*_AM_ > 300 in the conditions modeled in fig. 3B) to result in stable coexistence of two species. When that strong but incomplete, choice-based assortative mating induces low fitness of hybrids due to rare-mating-type disadvantage, amounting to substantial loss of hybrid fitness (e.g., a 30% loss; fig. 5). Hence when strong assortative mating does result in the maintenance of two species, it does so through impacts on both prezygotic and postzygotic isolation.

The effect of assortative mating on the fitness of mating types is dependent on their relative abundances, making systems dependent on their initial state. When *S*_AM_ = 1000 and *w*_hyb_ = 1, starting with two species results in maintenance of two species (fig. 5A), whereas starting with one intermediate species with substantial genetic variation would not result in the emergence of two species. At more moderate levels of assortative mating, the system can exhibit apparent stability as two species for a period of time, but then undergo a quick phase transition to a single species (fig. 5B).

The present simulations focus on cases of complete sympatry, and we can consider them in comparison with related simulations of hybrid zones between initially allopatric species in which individuals have limited dispersal distances (Irwin 2020). In the spatially structured case, a small reduction in hybrid fitness compared to the initial species causes a narrow and stable hybrid zone, preventing blending of the two species. Assortative mating is ineffective in preventing blending unless it is extremely strong, in which case it induces low hybrid fitness through rare-mating-type disadvantage (Irwin 2020). In the present models of pure sympatry, low hybrid fitness tends to lead to extinction (of one or both species), and strong assortative mating is needed for coexistence. Together, the two sets of models indicate that: 1) a small reduction in hybrid fitness can stabilize the presence of differentiated geographic forms; 2) low to moderate levels of assortative mating (up to 10x to 300x, depending on other parameters) are ineffective in preventing blending of two species; 3) strong assortative mating is needed for sympatric coexistence; 4) strong assortative mating between two species can itself cause low hybrid fitness; 5) other forms of low hybrid fitness can be stabilizing, reducing the level of assortative mating needed for stable coexistence between species; and 6) very low hybrid fitness in the absence of complete assortative mating is destabilizing. Although the present analysis focuses on coexistence outcomes when there is complete sympatry with no spatial structure, we note that limited spatial structure is known to enable a broader range of conditions under which two differentiated populations can persist on the landscape (Barton and Hewitt 1989; Irwin 2020); in most of those cases, however, the ranges are best considered parapatric rather than qualifying as true sympatric coexistence.

A widely observed biogeographic pattern is that the most closely related species pairs tend to be geographically separated, whereas sympatrically occurring species tend to be more distantly related (Weir and Price 2011), implying that insufficient differentiation in some way limits coexistence. After considering our simulation results, we postulate that achieving sympatry between related species tends to be limited more often by insufficient assortative mating than by insufficient ecological differentiation. Our reasoning is that sympatric coexistence is possible with just a little bit of ecological differentiation (fig. 6) but requires strong assortative mating under all modelled scenarios (e.g., at least 10x or more, depending on other parameters). The widespread observation of narrow hybrid zones that appear to prevent sympatric overlap of ranges of related species (Hewitt 1988; Barton and Hewitt 1989; Gompert et al. 2017) reinforces this view that hybridization is the primary limit on coexistence.

However, an important lesson of the simulations is that the effects on coexistence of incomplete assortative mating and insufficient ecological differentiation depend on the intrinsic growth rate (*R*; fig. 6). If *R* is high, then only a little bit of ecological differentiation (i.e., a small *E*) is needed to enable coexistence, and this can be maintained at a wider range of parameter values for assortative mating and hybrid fitness. If *R* is low, then more ecological differentiation is needed, and the range of parameter values is more restrictive. This important role for intrinsic growth rate points to the possibility of different taxonomic groups requiring different levels of assortative mating and ecological differentiation for stable coexistence. For instance, birds and mammals tend to have small clutch or litter sizes, meaning small *R* and limited ability to coexist without very strong assortative mating. In contrast, many groups such as insects, fish, and plants can have large numbers of offspring, potentially meaning high *R* and a broader range of conditions over which stable coexistence is possible.

These results indicate that the potential for cross-mating behavior and hybridization needs to be incorporated into coexistence theory. A recent review also advocated for between-species reproductive interactions being incorporated into coexistence theory (Gómez-Llano et al. 2021), although it emphasized how these interactions might facilitate coexistence between ecologically similar species. If we add the buildup of hybrids to this conceptual framework, the potential for coexistence of two species is reduced. Here, under most of the simulations presented (all those with complete ecological differentiation, where *E* = 1), the two species exploit different ecological niches and hence coexistence is predicted under ecological coexistence theory (Vandermeer 1972; Chesson 2000; Siepielski and McPeek 2010; HilleRisLambers et al. 2012; Mittelbach and McGill 2019). A small rate of hybridization can shift the outcome towards blending, possibly resulting in extinction of one or both species. The levels of hybridization at which such outcomes are observed might be difficult to detect when the two species are not yet sympatric: for example, many studies of mating behavior would not have the power to distinguish complete premating isolation from a 1/300 probability of a female choosing a heterospecific mate compared to a conspecific mate. Yet the long-term outcome of that level of interbreeding can be complete blending or extinction of one species (depending on other parameters). While it might seem that this need to incorporate hybridization into coexistence theory only applies to very closely related species, hybridization has been observed between species separated by many millions of years of evolution (Rothfels et al. 2015; Toews et al. 2020). Moreover, the simulations show that even when hybrids are inviable (*w*_hyb_ = 0), incomplete assortative mating can lead to failure of the two species to coexist. Hence even when hybrids are never observed (because zygotes do not develop), any tendency to interbreed between species can limit sympatric coexistence.

## Acknowledgements

For comments on the manuscript we thank Rachel Germain, Jessica Irwin, James Mallet, Ellen Nikelski, Jeannette Whitton, and members of the Irwin lab. We thank Jennifer Lau, Roger Butlin and two anonymous reviewers for comments during the review process. This research was supported by a grant from the Natural Sciences and Engineering Research Council of Canada (RGPIN-2017-03919 and RGPAS-2017-507830 to DEI).

## Data and Code Availability

The HZAM-Sym code for running simulations using R is available at https://github.com/darreni/HZAM-Sym, and the code using Julia is available at https://github.com/darreni/HZAM_Sym_Julia.

Data files, results graphs, and code are archived at Dryad: https://doi.org/10.5061/dryad.9ghx3ffhb

## Appendix Group-based assortative mating

As discussed in the Results section, the choice-based assortative mating used in the main analysis can induce sexual selection against rare mating types (fig. 5). We also explore a mating system that is group-based with random mating within each group, such that it avoids sexual selection. We think that choice-based assortative mating is generally more realistic than the group-based system, and we are including the methods and results from the latter here for those interested.

In the group-based method, individual females and males are assigned to mating groups based on their functional trait *T*, but with modified probabilities according a parameter σ_G_. Based on *L* genetic loci underlying the trait *T*, there are *g* = 2*L*+1 possible values for trait *T*, with values evenly spaced from zero to one, and hence *g* mating groups with each group corresponding to a primary value of *T*, *T*_G_. To determine which mating group each individual joins, we modify its trait *T* by adding a value drawn from a normal distribution with mean zero and standard deviation σ_G_. If the new modified value, *T*_m_, is outside of the range {(0 – 0.5/(*g*-1)), (1 + 0.5/(*g*-1)}, then that is rejected and a new draw is made, until *T*_m_ is within that range. Then, that individual is assigned to the group with the closest *T*_G_ value to the individual’s value of *T*_m_.

Once all individuals are in mating groups, mating pairs are determined randomly within each mating group, with each male and each female mating at most once with one other individual. This means that some individuals are left unpaired and do not reproduce. Due to the role of stochasticity in being higher in a small sample size, the probability of being unpaired tends to be higher when there are fewer individuals within the mating group (because the number of females and males are likely to be proportionally more different). To correct for this effect (which otherwise would introduce another form of sexual selection on the trait, due to group membership), we determine the fraction of individuals that are paired in a mating group, *F*_paired_, and adjust the expected number of offspring for each female (*c*) in the mating group by dividing by *F*_paired_.

To compare results from the choice-based mating system and those from the group-based mating system, we measured the fraction of offspring who are F1 hybrids produced by a single generation of reproduction at the start of each simulation, when the two species first become sympatric. This was done first for the choice-based model, measuring the fraction of F1 hybrids for various levels of *S*_AM_. Then, the group-based model was run at varying levels of σ_G_, tuning the values that produced equivalent proportions of F1 hybrids. Once these values of σ_G_ that give equivalent levels of interbreeding as the values of *S*_AM_ in the choice-based model, simulations were run for 1000 generations under similar parameter combinations as in the choice-based model.

**Figure A1.**
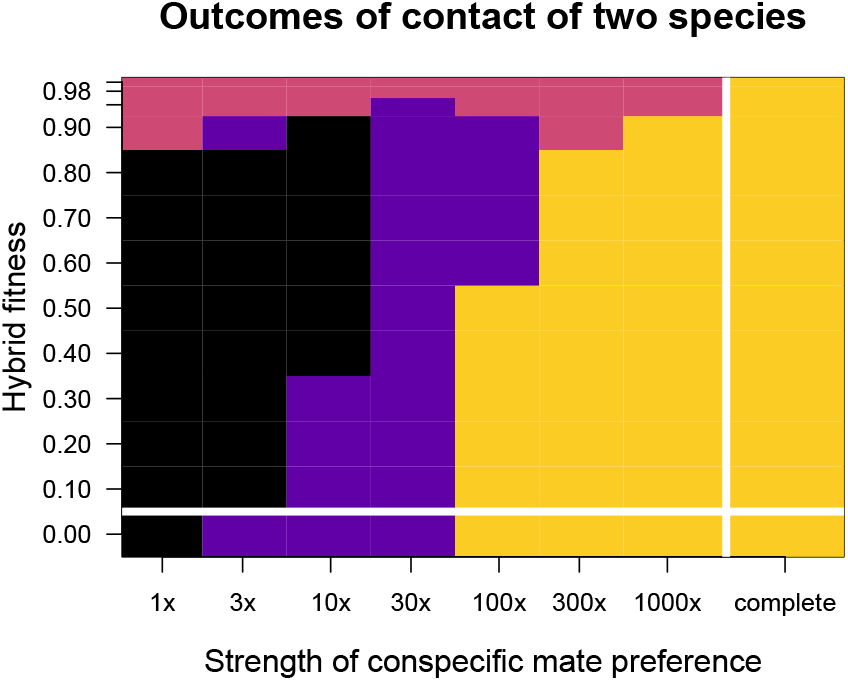
Outcomes of contact between two populations after 1000 generations in the group mating model, at various combinations of assortative mating strength (along horizontal axis) and hybrid fitness (vertical axis). Five simulations were run under each set of parameter combinations (as indicated by the marks along each axis), with colors representing the most common outcome for each set: black = extinction of both species; purple = one species remaining (extinction of the other); salmon = hybrid population; yellow = two species. In these simulations total carrying capacity is 1000 (500 on each of two resources) and growth rate *R* = 1.05. Compare to figure 3B for results under the same settings except with choice-based mating rather than group-based mating. Produced using the R implementation of HZAM.

Results of the group-based model are summarized in figure A1, which was produced under equivalent sets of parameter values as figure 3B, but with the group-based mating rather than choice-based mating.

When there is no explicit lower fitness of hybrids (i.e., the top row of results in fig. A1), any hybridization at all results in blending of the two species into a mixed population. This differs from the choice-based model, wherein strong choice-based assortative mating leads to low fitness of hybrids due to rare-mating-type disadvantage, stabilizing the presence of two species.

Another difference between results of the group-based model and the choice-based model is the parameter space over which extinction of both populations occurs, when hybrids are designated as having low intrinsic fitness. This is due to the choice-based model having a quicker evolution from a hybrid population to a single-species population, due to choice-based selection against rare types. Without this choice-induced selection, the group-based model stays longer in a mixed population, more often declining to extinction rather than recovering to one or the other species.

## References

Barton, N. H., and K. S. Gale. 1993. Genetic analysis of hybrid zones. Pages 13–45 *in* R. G. Harrison, ed. Hybrid zones and the evolutionary process. Oxford University Press, Oxford.

Barton, N. H., and G. M. Hewitt. 1989. Adaptation, speciation and hybrid zones. Nature 341:497–503.

Bezanson, J., A. Edelman, S. Karpinski, and V. B. Shah. 2017. Julia: a fresh approach to numerical computing. SIAM Review 59:65–98.

Chesson, P. 2000. Mechanisms of maintence of species diversity. Annual Review of Ecology and Systematics 31:343–66.

Cronemberger, Á. A., A. Aleixo, E. K. Mikkelsen, and J. T. Weir. 2020. Postzygotic isolation drives genomic speciation between highly cryptic *Hypocnemis* antbirds from Amazonia. Evolution 74:2512–2525.

De Cara, M. A. R., N. H. Barton, and M. Kirkpatrick. 2008. A model for the evolution of assortative mating. American Naturalist 171:580–596.

Durrett, R., L. Buttel, and R. Harrison. 2000. Spatial models for hybrid zones. Heredity 84:9–19.

Felsenstein, J. 1981. Skepticism towards Santa Rosalia, or Why are there so few kinds of animals? Evolution 35:124.

Garnier, S. 2018. viridis: default color maps from “matplotlib”. R package version 0.5.1.

Germain, R. M., S. P. Hart, M. M. Turcotte, S. P. Otto, J. Sakarchi, J. Rolland, T. Usui, et al. 2020. On the origin of coexisting species. Trends in Ecology & Evolution 1–10.

Germain, R. M., J. T. Weir, and B. Gilbert. 2016. Species coexistence: Macroevolutionary relationships and the contingency of historical interactions. Proceedings of the Royal Society B: Biological Sciences 283.

Gómez-Llano, M., R. M. Germain, D. Kyogoku, M. A. McPeek, and A. M. Siepielski. 2021. When ecology fails: how reproductive interactions promote species coexistence. Trends in Ecology & Evolution 1–13.

Gompert, Z., E. G. Mandeville, and C. A. Buerkle. 2017. Analysis of population genomic data from hybrid zones. Annual Review of Ecology, Evolution, and Systematics 48:207–229.

Haldane, J. B. S. 1948. The theory of a cline. Journal of Genetics 48:277–284.

Hewitt, G. M. 1988. Hybrid zones – natural laboratories for evolutionary studies. Trends in Ecology and Evolution 3:158–167.

HilleRisLambers, J., P. B. Adler, W. S. Harpole, J. M. Levine, and M. M. Mayfield. 2012. Rethinking community assembly through the lens of coexistence theory. Annual Review of Ecology, Evolution, and Systematics 43:227–248.

Irwin, D. E. 2020. Assortative mating in hybrid zones is remarkably ineffective in promoting speciation. The American Naturalist 195:E150–E167.

Kuno, E. 1992. Competitive exclusion through reproductive interference. Researches on Population Ecology 34:275–284.

Liou, L. W., and T. D. Price. 1994. Speciation by reinforcement of premating isolation. Evolution 48:1451–1459.

Mallet, J. 2005. Hybridization as an invasion of the genome. Trends in Ecology and Evolution 20:229–237.

Mikkelsen, E. K., and D. Irwin. 2021. Ongoing production of low-fitness hybrids limits range overlap between divergent cryptic species. Molecular Ecology 30:4090–4102.

Mittelbach, G. G., and B. J. McGill. 2019. Species coexistence and niche theory. Pages 141–157 *in* Community Ecology. Oxford University Press, Oxford, UK.

Otto, S. P., M. R. Servedio, and S. L. Nuismer. 2008. Frequency-dependent selection and the evolution of assortative mating. Genetics 179:2091–2112.

Panhuis, T. M., R. Butlin, M. Zuk, and T. Tregenza. 2001. Sexual selection and speciation. Trends in Ecology & Evolution 16:364–371.

Pennings, P. S., M. Kopp, G. Meszéna, U. Dieckmann, and J. Hermisson. 2008. An analytically tractable model for competitive speciation. American Naturalist 171.

Perini, S., M. Rafajlović, A. M. Westram, K. Johannesson, and R. K. Butlin. 2020. Assortative mating, sexual selection, and their consequences for gene flow in *Littorina*. Evolution 74:1482–1497.

Polechová, J., and N. Barton. 2011. Genetic drift widens the expected cline but narrows the expected cline width. Genetics 189:227–235.

Price, T. 2008. Speciation in Birds. Roberts & Company Publishers, Greenwood Village, Colorado.

Prout, T. 1978. The joint effects of the release of sterile males and immigration of fertilized females on a density regulated population. Theoretical Population Biology 13:40–71.

Pulido-Santacruz, P., A. Aleixo, and J. T. Weir. 2018. Morphologically cryptic Amazonian bird species pairs exhibit strong postzygotic reproductive isolation. Proceedings of the Royal Society B: Biological Sciences 285:20172081.

R Core Team. 2021. R: a language and environment for statistical computing. R Foundation for Statistical Computing, Vienna, Austria.

Rothfels, C. J., A. K. Johnson, P. H. Hovenkamp, D. L. Swofford, H. C. Roskam, C. R. Fraser-Jenkins, M. D. Windham, et al. 2015. Natural hybridization between genera that diverged from each other approximately 60 million years ago. American Naturalist 185:433–442.

Schluter, D., and M. W. Pennell. 2017. Speciation gradients and the distribution of biodiversity. Nature 546:48–55.

Servedio, M. R., and J. Hermisson. 2020. The evolution of partial reproductive isolation as an adaptive optimum. Evolution 74:4–14.

Siepielski, A. M., and M. A. McPeek. 2010. On the evidence for species coexistence: a critique of the coexistence program. Ecology 91:3153–3164.

Singhal, S., and C. Moritz. 2012. Strong selection against hybrids maintains a narrow contact zone between morphologically cryptic lineages in a rainforest lizard. Evolution 66:1474–1489.

Taylor, S. A., and E. L. Larson. 2019. Insights from genomes into the evolutionary importance and prevalence of hybridization in nature. Nature Ecology and Evolution 3:170–177.

Toews, D. P. L., G. R. Kramer, A. W. Jones, C. L. Brennan, B. E. Cloud, D. E. Andersen, I. J. Lovette, et al. 2020. Genomic identification of intergeneric hybrids in New World wood-warblers (Aves: Parulidae). Biological Journal of the Linnean Society 131:183–191.

Turelli, M., N. H. Barton, and J. A. Coyne. 2001. Theory and speciation. Trends in Ecology & Evolution 16:330–343.

Vandermeer, J. H. 1972. Niche theory. Annual Review of Ecology and Systematics 3:107–132.

Weir, J. T., and T. D. Price. 2011. Limits to speciation inferred from times to secondary sympatry and ages of hybridizing species along a latitudinal gradient. American Naturalist 177:462–469.

Whitton, J., C. J. Sears, and W. P. Maddison. 2017. Co-occurrence of related asexual, but not sexual, lineages suggests that reproductive interference limits coexistence. Proceedings of the Royal Society B: Biological Sciences 284:20171579.

