## Supplementary figures for "Hybridization and the Coexistence of Species"

All simulations and graphs shown in this supplement were produced using the Julia implementation of HZAM-Sym.

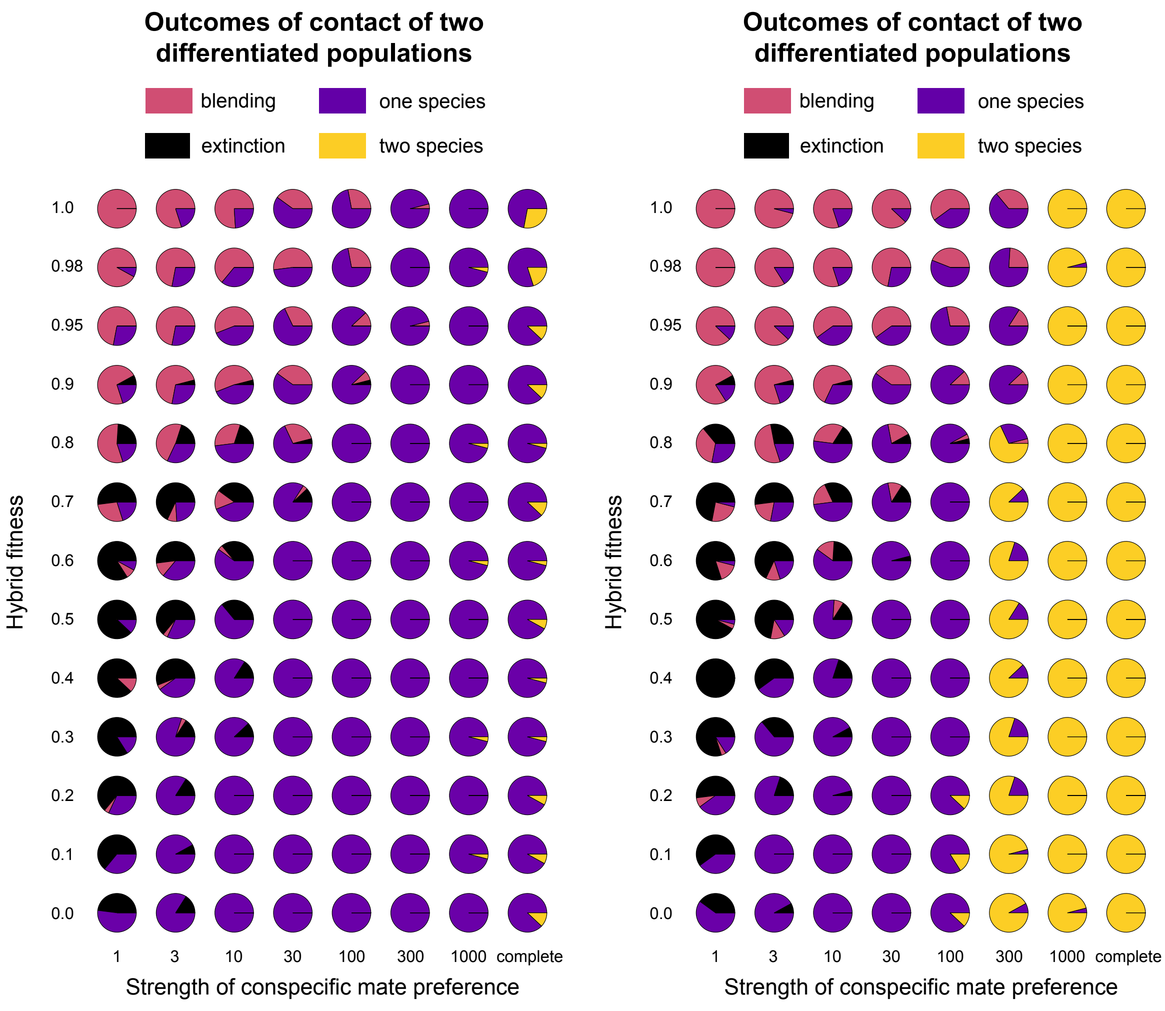


**Figure S1.** Frequencies of four outcomes under each set of conditions summarized in figure 3 (which showed just the most common outcome for each parameter set). Each pie graph shows the frequency of outcomes among 25 replicate simulations under that set of conditions. The left panel corresponds to figure 3A (no ecological differentiation), and the right to figure 3B (full ecological differentiation).

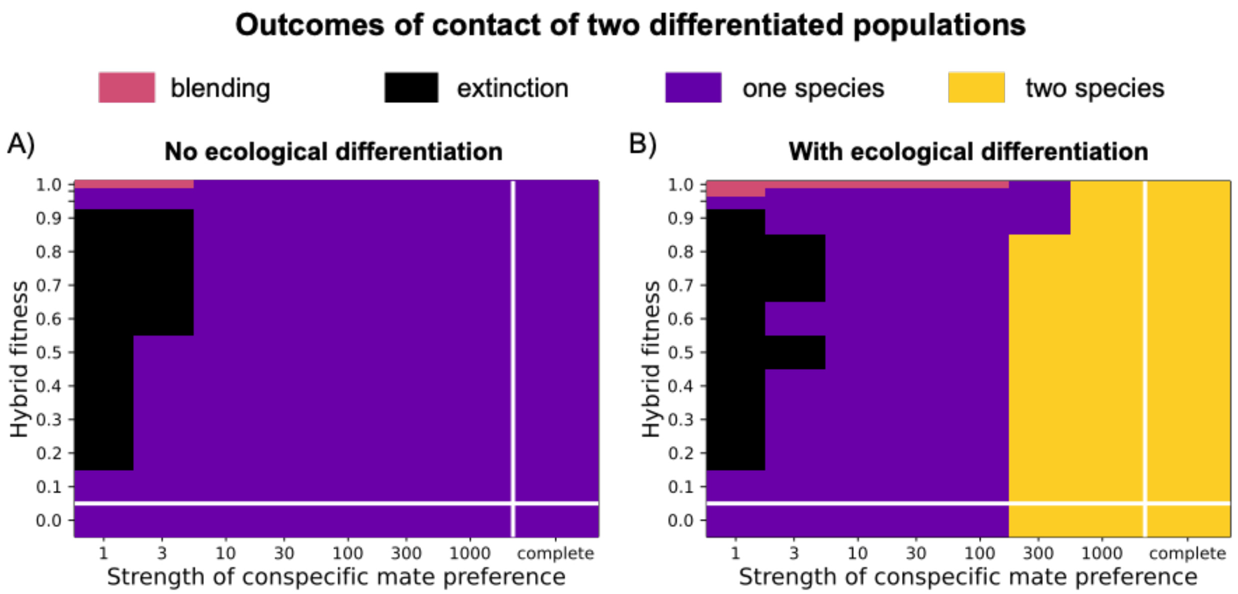


**Figure S2.** Most frequent outcomes for simulations conducted under identical conditions to figure 3, but with epistasis-based survival fitness rather than underdominance-based. There are 25 replicates under each parameter set (5200 simulations total for this figure); see figure S3 for the detailed breakdown of frequencies of outcomes for each parameter set.

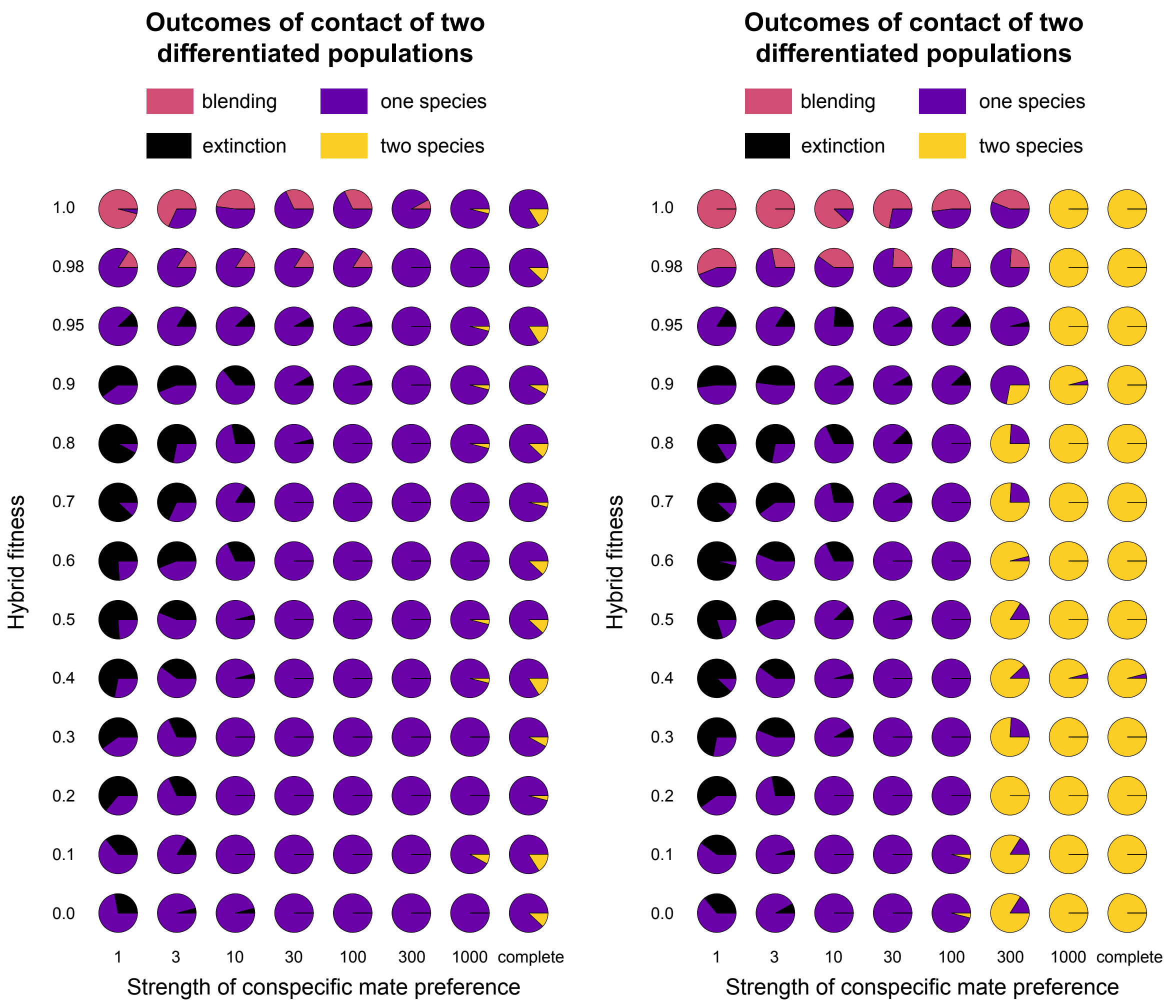


**Figure S3.** Frequencies of four outcomes for the 5200 simulations summarized in figure S2.

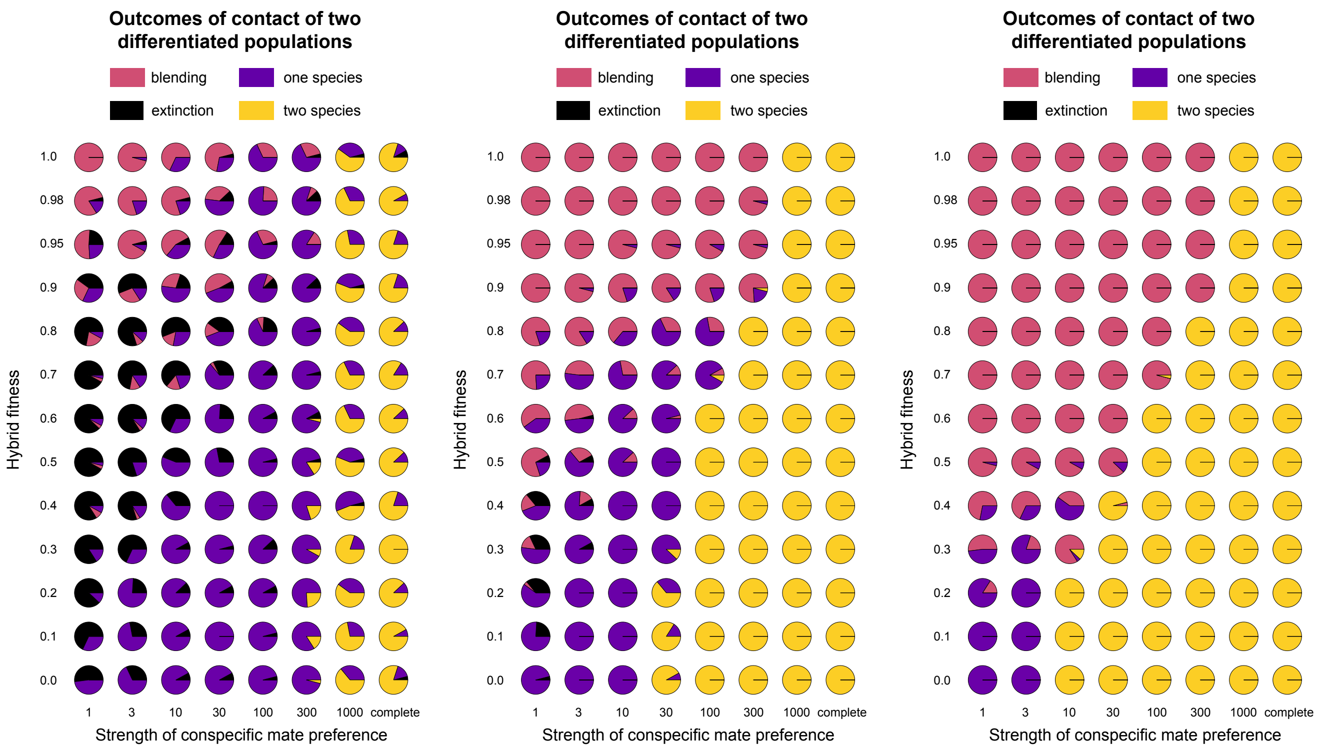


**Figure S4.** Frequencies of four outcomes for the 7800 simulations summarized in figure 4 (see caption of that figure for details). Left: R = 1.025. Middle: R = 1.2. Right: R= 2.6.

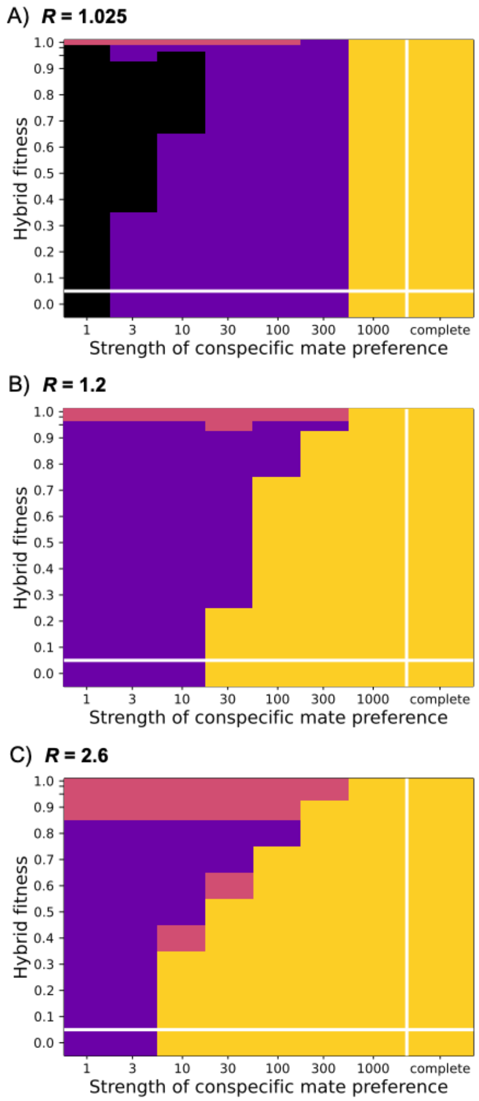


**Figure S5.** Most frequent outcomes for simulations conducted under identical conditions to figure 4, but with epistasis-based (rather than underdominance-based) survival fitness. There are 25 replicates under each parameter set (7800 simulations total for this figure); see figure S5 for a detailed breakdown of frequencies of outcomes for each parameter set.

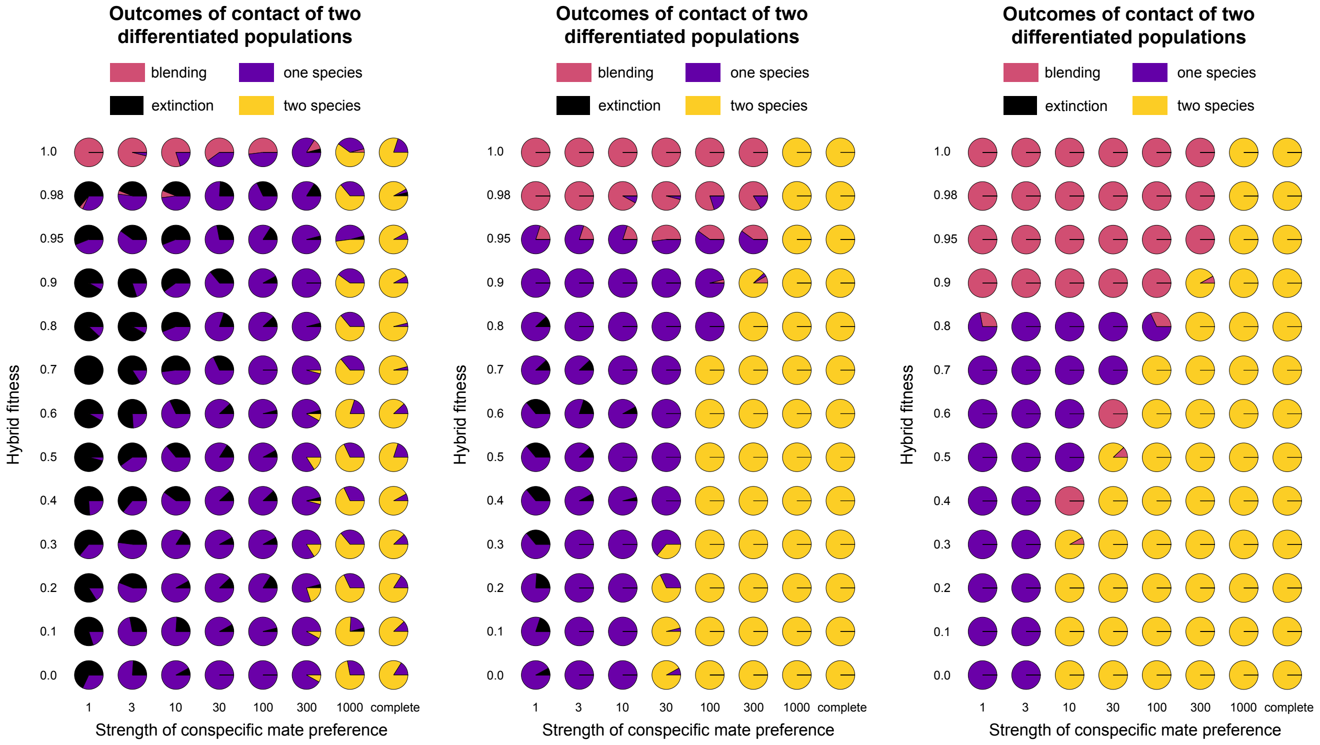


**Figure S6.** Frequencies of four outcomes for the 7800 simulations summarized in figure S5 (see caption of that figure for details). Left: R = 1.025. Middle: R = 1.2. Right: R= 2.6.

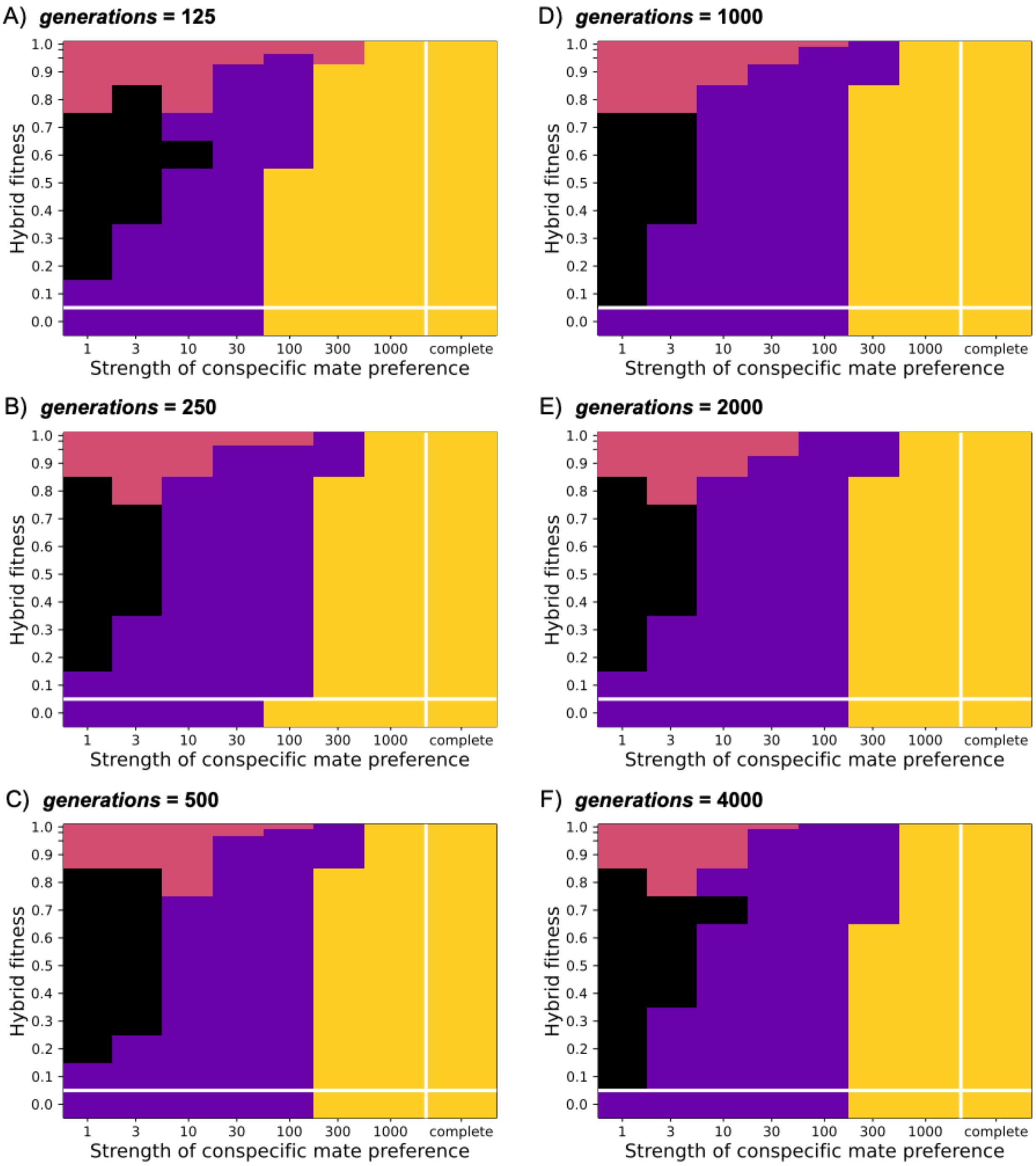


**Figure S7.** Comparison of how coexistence outcomes relate to the number of generations the simulations run. These simulations were run under identical settings as figure 3B, but for varying number of generations as indicated above each panel (ranging from 125 in A to 4000 in F). Figure 3B is identical to panel D of this figure. Twenty-five simulations were run for each combination of parameters (a total of 15600 for this figure), with the most common outcome shown for each (see caption of figure 3 for the key to colors of outcomes.)

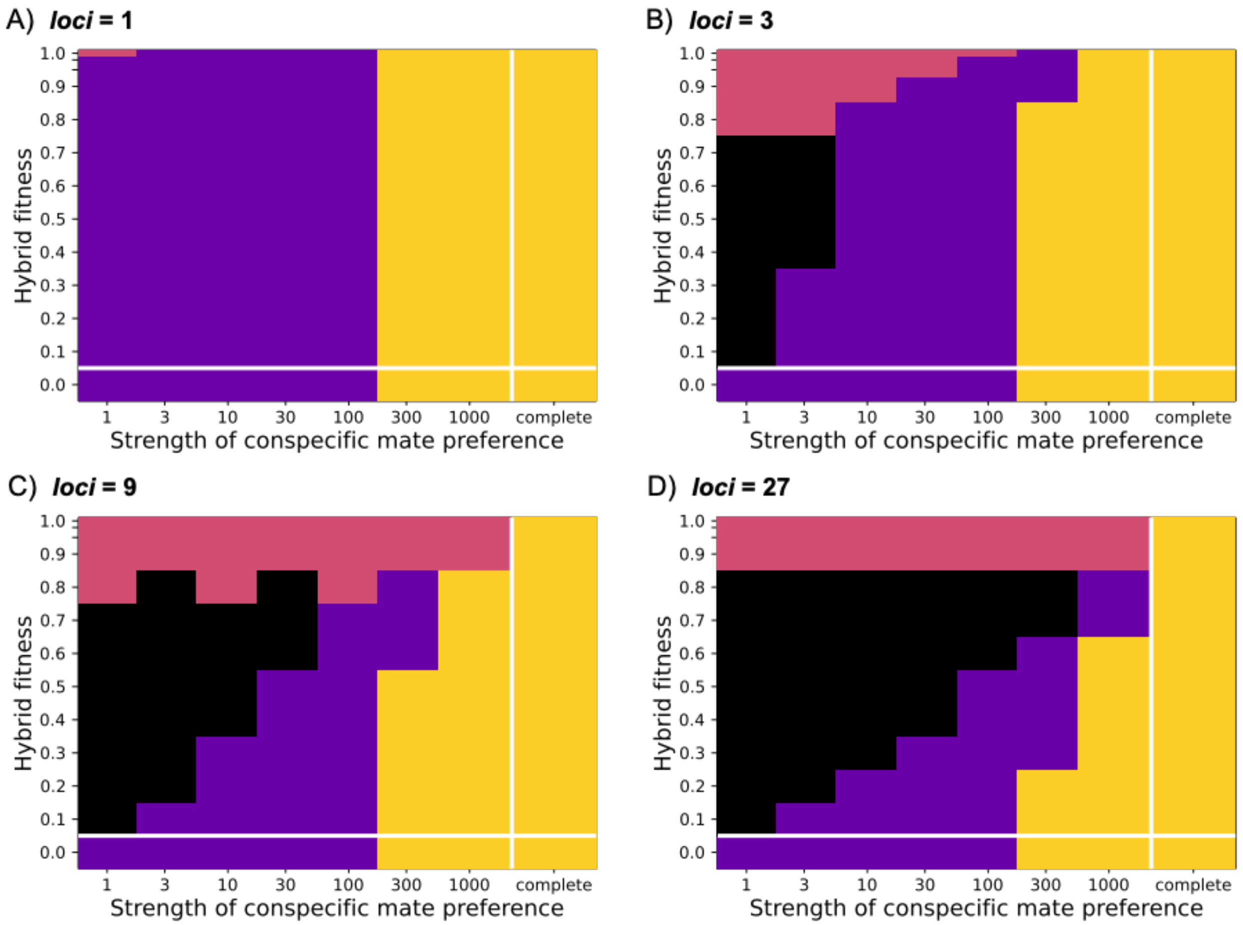


**Figure S8.** Examination of how coexistence outcomes relate to the number of loci. These simulations were run under identical settings as figure 3B, but for varying numbers of loci, as indicate above each panel (ranging from 1 in A to 27 in D). Twenty-five simulations were run for each combination of parameters (so 10400 simulations for this figure), with the most common outcome shown for each. Figure 3B is identical to panel B of this figure.

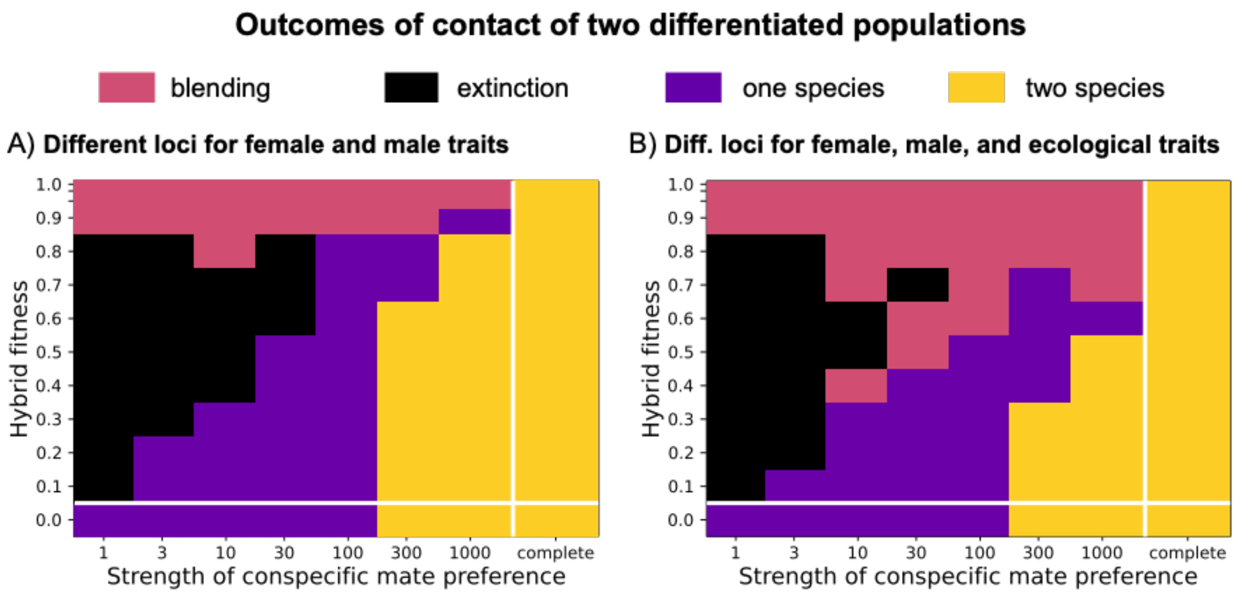


**Figure S9.** The most common outcomes for conditions identical to figure 3B except with (A) different loci for female and male traits (3 of each, for a total of 6), or (B) different loci for female traits, male traits, and ecological traits (3 of each, for a total of 9). Twenty-five simulations were run for each combination of parameters (so 5200 total), with the most common outcome shown for each parameter combination.

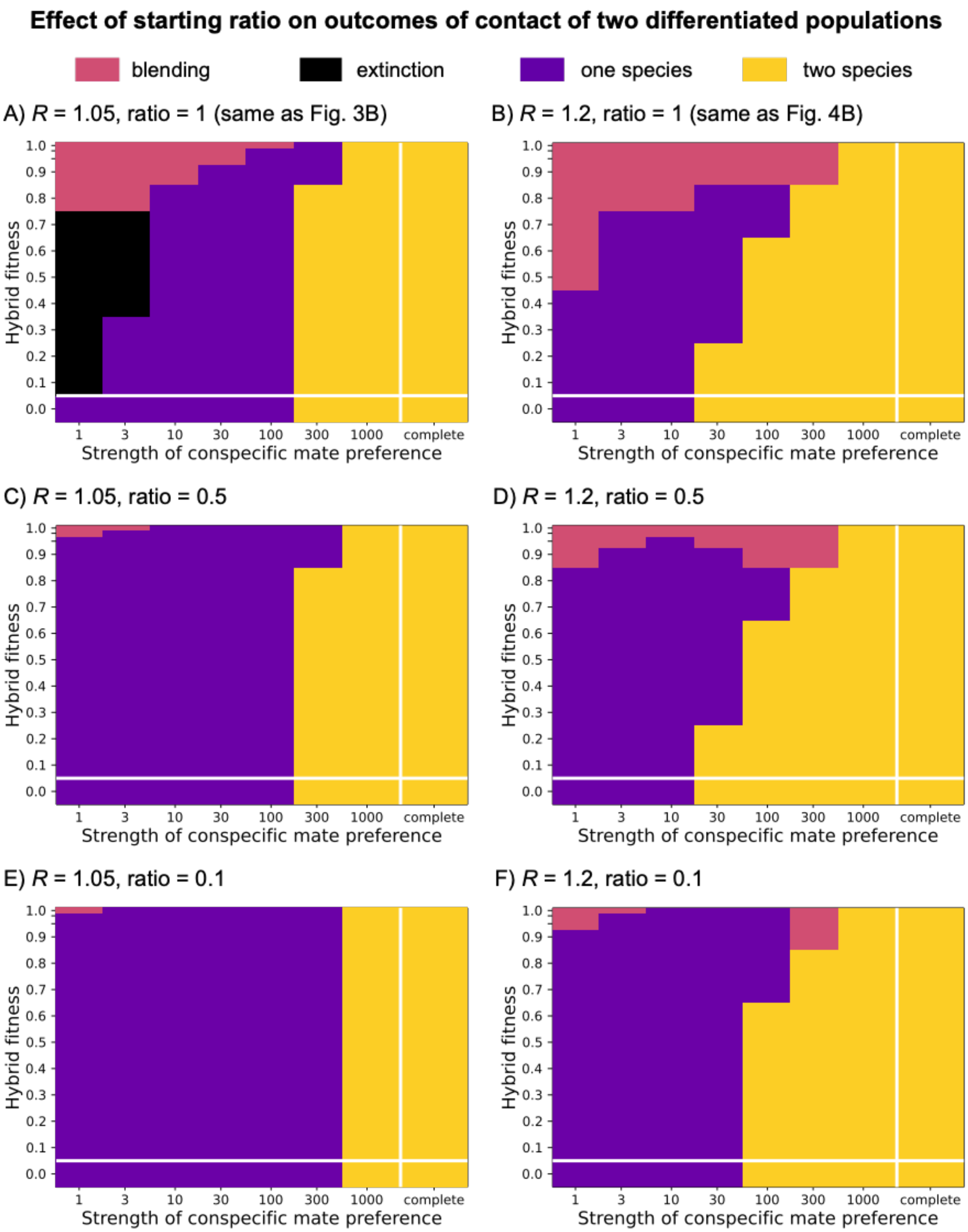


**Figure S10.** Examination of how ratio of initial population sizes affects sympatric coexistence outcomes. The left column is for $R$ = 1.05, and the right for $R$ = 1.2. The top two panels (A, B) are outcomes for equal starting sizes of the two populations, whereas the middle row (C, D) are outcomes when one starting population is 0.5 the size of the other, and the lower row (E, F) are outcomes when one starting population is 0.1 the size of the other. All other settings are identical to those of figures 3B and 4B.
